# Three-dimensional interactions between integrated HPV genomes and cellular chromatin dysregulate host gene expression in early cervical carcinogenesis

**DOI:** 10.1101/2021.02.03.429496

**Authors:** Ian J Groves, Emma LA Drane, Marco Michalski, Jack M Monahan, Cinzia G Scarpini, Stephen P Smith, Giovanni Bussotti, Csilla Várnai, Stefan Schoenfelder, Peter Fraser, Anton J Enright, Nicholas Coleman

## Abstract

Development of cervical cancer is directly associated with integration of human papillomavirus (HPV) genomes into host chromosomes and subsequent modulation of HPV oncogene expression, which correlates with multi-layered epigenetic changes at the integrated HPV genomes. However, the process of integration itself and dysregulation of host gene expression at sites of integration in our model of HPV16 integrant clone natural selection has remained enigmatic. We now show, using a state-of-the-art ‘HPV integrated site capture’ (HISC) technique, that integration likely occurs through microhomology-mediated repair (MHMR) mechanisms via either a direct process, resulting in host sequence deletion (in our case, partially homozygously) or via a ‘looping’ mechanism by which flanking host regions become amplified. Furthermore, using our ‘HPV16-specific Region Capture Hi-C’ technique, we have determined that three-dimensional (3D) interactions between the integrated virus genome and host chromosomes, both at short- (<500 kbp) and long-range (>500 kbp), appear to drive host gene dysregulation through the disruption of local host:host 3D interactions known as topologically associating domains (TADs). This mechanism of HPV-induced host gene expression modulation indicates that integration of virus genomes near to or within a ‘cancer-causing gene’ is not essential to influence such genes within an entire TAD and that these modifications to 3D interactions could have a major role in selection of HPV integrants at the early stage of cervical neoplastic progression.

## Introduction

Human papillomavirus (HPV) infection is associated with the development of around 5% of all human cancers, with ~690,000 cases arising annually worldwide(1). Of these, ~570,000 are cancers of the cervix, which usually present as squamous cell carcinomas (SCCs), developing through clonal selection of cells with a competitive growth advantage from pre-cursor squamous intraepithelial lesions (SILs) and ultimately leading to ~260,000 deaths globally(2-5). The association of high-risk HPV (HRHPV) infections with cervical SCCs is over 99.9% and, as such, makes HPV the etiological agent associated with cervical carcinomas(6, 7). The treatment of HPV-associated carcinomas has changed little over the past 30 years and, despite current vaccination programs against HPV, new therapies are necessary for an aging unvaccinated population.

The genome of HPV usually exists as an extra-chromosomal episome of around 7.9 kilobases (kb) at a copy number of around 100 per cell in squamous epithelial lesions as part of the normal virus lifecycle(8, 9). Although development of cervical SCC with an episomal HPV genome can occur(10), progression of disease toward cervical carcinoma is more often associated with integration of the fractured double stranded DNA (dsDNA) HRHPV genome into that of the host, occurring in around 85% of cases(11-13). The integration process usually involves the disruption of the HPV *E2* gene and, with loss of this trans-repressive protein product, leads to dysregulation of virus gene expression from the early promoter(5, 11, 14). Despite this process usually resulting in an increase in HPV oncogene expression associated with cervical SCCs, our previous work has shown that the genomes at initial integration events prior to selection can have levels of expression similar to, or lower than, episomal parental cell lines(15).

The control of HPV gene expression in the productive lifecycle of the virus occurs in a differentiation-dependent manner associated with the position of the virus within the infected stratified epithelium(5). The necessary expression of the HPV oncogenes *E6* and *E7* in the basal layer cells occurs through transcriptional initiation at the virus early promoter (p97 in the case of HPV16). This is controlled by the interaction of various host transcription factors with regulatory elements within the long control region (LCR) and modification to the local chromatin structure(5, 16, 17). The binding of these factors is known to become modified as infected cells differentiate such that the late promoter (p670 for HPV16) is stimulated and late virus gene expression ensues(18, 19). In concert, changes to chromatin structure are known to occur as late HPV gene expression becomes activated, driven through the modification of histone post-translational modifications (PTMs) at HPV genome-associated nucleosomes(18, 20). These histone PTMs have also been found associated with the enhancer and promoters of HPV16 episomes during progression of *in vitro* neoplastic progression with acetylation of both histone H3 and H4 (H3ac and H4ac, respectively) accumulating during the initial stages of phenotypic progression to SCC(10). Work from our lab investigating the integrated HPV16 genome has previously shown differential association of histone modifications with the virus LCR and early genes corresponding to levels of virus transcript per template(15). Subsequent studies have shown that multi-layered epigenetic modifications to the integrated HPV genome are associated with the recruitment and activation of RNA polymerase II (RNAPII), thereby determining the level to which HPV oncoprotein expression occurs(21). These modifications include levels of DNA methylation, further histone PTMs and associated enzymes as well as nucleosome positioning and the presence of chromatin remodelling enzymes and transcriptional activators, such as the P-TEFb complex, directly at the integrated virus templates(5, 21).

However, as implied, cervical SCC is not always associated with high virus oncogene levels and may be driven independent of this factor, for example through host gene changes(22). HPV appears to integrate into certain sites across the human genome more often than others, so called ‘integration hotspots’, associated at times with chromosome fragile sites (CFSs)(12, 23-25). Integration can occur directly into a gene, both introns and exons, and can lead to varying changes in gene expression level(26-28). In a study of HPV-positive head and neck SCC (HNSCC), 17% of integrants were also found within 20 kbp of a gene, indicative of possible selective pressure from integration at, or near to, coding regions through modifications to host gene expression(29). Varying explanations for modified gene expression include HPV integration as being commonly associated with amplification of the local region or indeed rearrangement and translocation of that region elsewhere(27, 29-34). Other studies have suggested that higher level transcriptional control may be at play: integration into flanking regions of genes, sometimes as far away as 500kb, has been found associated with large increases in gene transcription, including at the *MYC* and *HMGA2* genes(26, 27). Interestingly, the association of the 8q24.21 region within which the *MYC* gene resides has been highlighted in several studies previously(35-37) and was more recently investigated using genome-wide studies. Using RNA-seq, haplotype resolved data showed that MYC is highly overexpressed from the HPV18 integrated allele (95:1), which is also associated with higher levels of transcriptionally active chromatin marks, transcription factors and RNAPII(38). The analysis of ‘chromatin interaction analysis with paired-end tag’ ChIA-PET sequencing data pointed toward a long-range *cis* interaction between the integrated HPV18 promoter/enhancer and the *MYC* gene(38). Hence, is likely that three-dimensional (3D) interactions between distant regions of chromatin have ultimately driven selection of this cell line from an excised lesion.

The cloned cell lines by which we have previously shown epigenetic control of virus early gene expression from integrated HPV16 genomes were developed from the W12 cell model(24, 39, 40). This model was generated by primary culture of a cervical low-grade SIL (LSIL) from which keratinocytes naturally infected with HPV16 were able to grow in monolayer maintaining the virus episome copy number between 100-200 copies per cell(8). From this polyclonal cell population, continuous *in vitro* passage of cells in long-term culture is associated with the loss of episomes alongside the outgrowth of cells containing integrated HPV16 genomes, mirroring phenotypic progression *in vivo* from LSIL to high-grade SIL (HSIL) to SCC(10, 41). We have previously used this approach to develop many series from the W12 cell system, including cloned cell lines that have a range of discrete integration sites, similar in position to those seen *in vivo* in cervical SCC, occurring through integration events prior to selection of a clone with the greatest growth advantage(5, 23, 24, 42, 43). This was conducted with single cell cloning of an early passage (p12 and p13) of W12 culture series-2 (W12Ser2), from which an integrant at chromosome position 8q24.21 had outgrown in previous studies, and under non-competitive conditions whereby a repressive environment for integrants was maintained until after cloning(44). Although the number of virus genomes associated with each cloned line varied, only a small number displayed concatemerisation of virus genomes (type II integrants) and, as such, the majority were deemed type I integrants displaying a premalignant LSIL-like phenotype when grown in organotypic ‘raft’ culture(4, 15). These clones therefore represent a typical population of polyclonal cells found in a premalignant cervical lesion and can be used to determine the factors that drive selection of certain integrants.

In work presented here, we have used the W12 clones to investigate further features of genome integration and host gene expression modulation that could lead to the selection of individual cells during carcinogenesis. Using cells that constitute type I integrants with four or less copies of integrated genomes that have expression per template levels of oncogenes that encompass the range seen within the total panel of W12 integrant clones(21), we have developed a novel and state-of-the-art technique (HPV integrated site capture/HISC) to determine HPV16 integration sites genome-wide at nucleotide resolution while utilising HPV16-specifc Region Capture Hi-C to investigate potential 3D interactions between the integrated virus genome and host chromatin. We have been able to pinpoint the precise locations of HPV16 integration sites within our W12 integrant clones, coinciding with areas of open chromatin, as well as determining that integration likely occurs through microhomology-mediated repair mechanisms. Integration occurs through either a direct process, whereby regions of the host genome are deleted – in our case partially homozygously – or via a ‘looping’ mechanism by which flanking regions of the host genome become amplified. Furthermore, application of our Region Capture Hi-C technology has determined that 3D loops do indeed exist between the integrated virus genome and host chromatin both at short- (<500 kbp) and long-range (>500 kbp). Alongside RNA-seq data, we have also confirmed that these interactions do appear to drive host gene dysregulation, possibly through the disruption of the normal nuclear architecture within topologically associating domains (TADs), leading to individual gene and cross-region changes in host gene expression during the early stages of cervical neoplastic progression.

## Results

### Integrated HPV16 genomes interact in three-dimensions (3D) with host chromosomes

As it had previously been shown that interactions between an integrated HPV genome and host chromatin on the same chromosome could lead to changes to host gene expression that could be selected for during carcinogenesis(38), we wished to determine whether these 3D interactions were occurring at an earlier stage of cervical neoplasia in our HPV16-positive W12 integrant clones prior to selection. To address this, we first developed a HPV16-specific Region Capture Hi-C protocol (Supplementary Figure 1A) similar to that published previously(45, 46), here using biotinylated RNA baits specific for the HPV16 genome that would select chimeric DNA complexes from a Hi-C library based upon the virus sequence and hence allow determination of any 3D interactions in our five integrant clones previously epigenetically characterized(21). The regions of significant interaction between the HPV16 and host chromatin were determined using GOTHiC software and visualised using the Circos tool (Figure 1). In W12 clone G2, reads originate from all MboI fragments across the HPV16 genome, indicating interactions with the host and, although the distribution of the reads was fairly uniform, the greatest percentage of reads came from HPV16 gene *E7* (Figure 1A). Significant interactions occurred exclusively with chromosome 5, the chromosome of integration, with the majority likely due to *cis* interactions with bordering host sequences at the breakpoint. However, upon closer inspection of the single chromosome view (Figure 1A inset), there was a divergence between the majority of reads from the virus — indicating the integration site — and a subset of reads that mapped to a separate region of the host, indicating a long-range 3D interaction between the integrated HPV16 genome in G2 and the host. In clone D2 (Figure 1B), the HPV16 genome also interacted with the site of integration at chromosome 5; however, the virus-host reads, predominantly from the *L2* gene, at this scale appear to converge on a single point at the host chromosome (Figure 1B inset). For three further W12 integrant clones (clones H, F and A5) interactions again occurred across the HPV16 genome with the host chromosome of integration, with the majority of reads coming from the *E2* portion of the virus genome for all (Figure 1C-E). In clone H, HPV16 integrated into chromosome 4 and resulted in a large deletion of the host (~170 kbp), which is illustrated by the separation of the virus-host reads in the chromosome view (Figure 1C inset). Interestingly, and contrary to our previous publications(15, 21, 24), we found that W12 clones F and A5 had the same integration site, with virus-host reads converging to the same region of chromosome 4 (Figure 1D & E).

**Figure 1.**
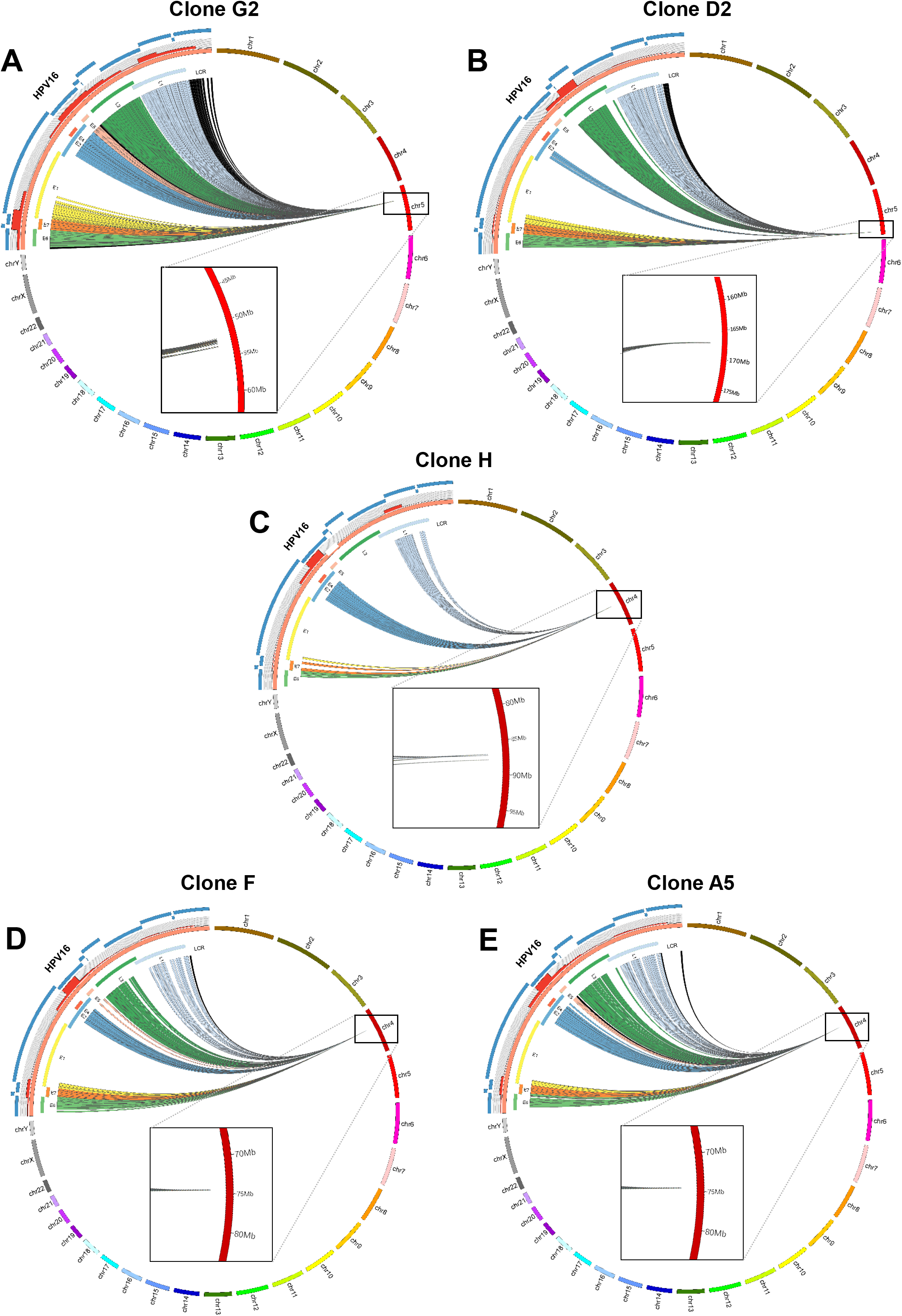
HPV16-specific Region Capture Hi-C determines definitive HPV16 integration sites. CIRCOS plots show sequence interactions between HPV16 (orange) and host chromosomes (various) for clones (A) G2, (B) D2, (C) H, (D) F and (E) A5. Each line within the circle represents a significant virus-host read indicating an above background interaction between a region of the HPV16 genome and the host. Reads are coloured to match individual HPV16 genes: E6 = green, E7 = orange, E1 = yellow, E2 = blue, E4 = red, E5 = pink, L1 = dark green, L2 = light blue and non-coding regions = black. Percentage of reads coming from different regions of the virus is indicated by the histogram on the outside of the HPV16 genome, which is split into 500 bp windows (red bars). HPV16 RNA bait fragments used in the Capture Hi-C experiment are indicated on the outside of the CIRCOS plot (blue curved lines). Presented data were generated using the Gothic program and plots are not to scale. Insets show zoomed sites of integration, with interaction divergence in clones G2 and H.

### HPV16 integration site virus-host breakpoint identification at nucleotide resolution

Having developed the probability that 3D virus-host interactions did indeed occur soon after HPV16 integration, we next sought to precisely identify the host sequence of the virus-host junctions. To do this with sufficient depth from genomic DNA samples, we developed a novel protocol by which DNA from each integrant clone was enriched for HPV16 sequences along with its flanking regions before sequencing, henceforth known as HPV Integration Site Capture (HISC) (Supplementary Figure 1B). The resulting sequencing data was analysed for reads mapping to the HPV16 genome with the corresponding human tag being determined. These data were then aligned to the HPV16 genome (Supplementary Figure 2) and the human genome (Supplementary Figures 3-6). From all integrant cell lines, sequencing reads mapped to both the HPV16 and host genomes with peaks at two distinct sites each (Supplementary Figures 2-6, HISC track), regardless of HPV16 genome copy number, demonstrating that only a single 5’ and 3’ breakpoint existed in each W12 clone examined. The separation of the breakpoint peaks from the HPV16 genome was consistent with termination of RNA-seq reads from separate transcriptome analysis (Supplementary Figure 2, RNA-seq tracks) and the known deletion of a proportion of the virus genome in each cell line(21), with the HPV16 transcription profiles additionally being consistent with our previously published quantitative PCR data(15). Alignment of the breakpoint peaks with host sequence gave a separation distance ranging from ~25-170 kbp across the integrant clones; the separation of peaks here however is consistent with two processes of HPV integration. The greatest distance of 170 kbp seen in clone H occurred due to the deletion of a proportion of the host genome upon ‘direct’ integration of the single HPV16 genome here determined by low coverage DNA-seq analysis (data not shown) and quantitative PCR (qPCR) of sections of host genome spanning the integration site (Supplementary Figure 5B). The breakpoint peak separation distance for the other four clones, however, was due to a ‘looping’ mechanism of HPV integration(47) whereby a flanking length of host sequence is amplified during integration of one or more copies of the virus. Again, this was verified through qPCR of host DNA across the integration sites (Supplementary Figures 3B, 4B & 6B), with transcription clearly occurring through these amplified host regions driven from the integrated HPV16 early promoter (Supplementary Figures 3-6A). The resultant genomic effect on the structure of both host alleles is pictured in Supplementary Figures 3-5C and 6D.

To confirm the virus-host breakpoints determined by HISC, PCR across each junction was carried out and Sanger sequenced with the verified coordinates of breakage summarized in Supplementary Figure 7. Interestingly, a 53 bp region of the *E2* gene was found to have been inverted after an intermediate 20 bp deletion, which had not been highlighted by alignment of RNA-seq reads to the HPV16 genome due to bioinformatics processing (Supplementary Figure 7D). Quantitative PCR (qPCR) was also carried to show that both the 5’ and 3’ breakpoints were unique to each cloned line (other than F and A5) with either very low or non-existent products produced with DNA from an early episomal population of W12 cells (W12par1 p12) or a normal cervix line, NCx/6 (Supplementary Figure 8).

Identification of the precise virus-host breakpoints in each cloned line allowed the extent of microhomology at integration sites to be assessed. At all integration sites, at least one end of the insertion involved nucleotides from both genomes directly adjacent to the junction being homologous, with a mode of 5 nt (range 3-5 nt) (Supplementary Figure 9A-D), with clone G2 having 5 nt homology at both ends (Supplementary Figure 9A). Additionally, when microhomology of 10 nt either side of each breakpoint was compared to that generated from 10,000 random shuffles of each sequence extended to 1,000 nt, 4 of 5 integrant clones (G2, H, F and A5) had statistically significant homology at one flank (Supplementary Figure 9A, C & D).

### HPV16 integrates into regions of open and transcriptionally active host chromatin

Mapping of the HPV16 integration sites from each cloned cell line allowed investigation of the genomic location and epigenetic landscape into which the virus had inserted (Supplementary Figure 10). In 4 of 5 cases, HPV16 had integrated into a gene (D2, TENM2; H, MAPK10; F & A5; RASSF6) with all of these cases arising at introns (Supplementary Figure 10B-D). Despite the integration site in clone G2 occurring intergenically, all insertions occurred at locations of open chromatin, characterized by DNaseI hypersensitivity sites from publicly available normal human epidermal keratinocyte (NHEK) ENCODE datasets (www.encodeproject.org). Additionally, the integration sites showed higher than average levels of histone post-translational modification marks associated with enhancers and transcriptional activity, namely H3K27ac and H3K4me1/2/3, with the marked exclusion of transcriptionally repressed facultative heterochromatin mark, H3K27me3. Noticeably, although integration in clone H did not occur directly into one of these loci, the HPV16 genome is located within 300 kbp of a similar site (Supplementary Figure 10C). Presence of binding sites of host architectural protein CTCF was found across the sites of integration, although again with higher than average occurrence, and usually within ~20 kbp of the inserted HPV16 genome.

### Short- and long-range 3D interactions occur between the HPV16 and host genomes

To inspect 3D interactions between the HPV16 and host genomes, Region Capture Hi-C sequencing data was re-visualised using SeqMonk software (Babraham Bioinformatics) and again aligned to NHEK ENCODE datasets at the location of integration for each cloned cell line. Local inspection of the points of 3D interaction with the host genome at the integration site of clone G2 showed highest levels corresponding to the 5’ and 3’ virus-host junctions (Figure 2A, red bars), with above background interactions occurring within this region of host genome amplification. This was found consistently across the Region Capture Hi-C datasets for all clones (Figure 3A & Supplementary Figure 11), with the exclusion of clone H where the absence of any intermediate interactions is likely due to deletion of this host region during integration (Supplementary Figure 11A).

**Figure 2.**
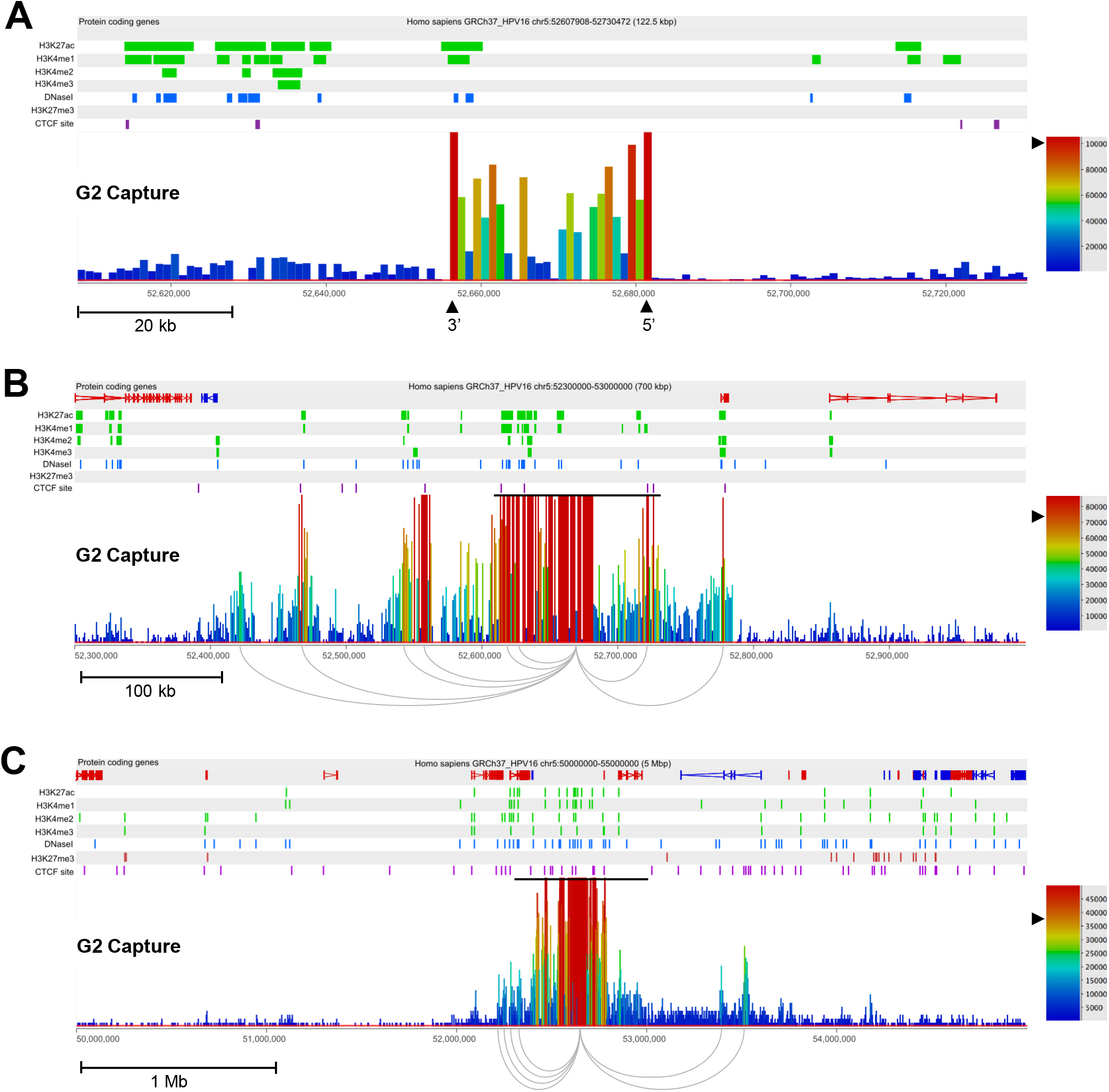
Identification of short- and long-range interactions between integrated HPV16 genomes and the host chromosome in W12 clone G2. (A) Capture Hi-C data is presented 122.5 kbp across the HPV16 integration locus. The 5’ and 3’ breakpoints of the virus are indicated by the tallest red bars and are labelled with black arrowheads, being inverted in comparison to the direction of host sequence due to the ‘looping’ integration mechanism. (B) Capture Hi-C data is presented 700 kbp across the HPV16 integration locus. The black line above the read peaks indicates the genomic window seen in panel A. Peaks of reads indicate regions of the host interacting with the integrated virus in three-dimensions. Short-range interactions between the HPV16 genome and host regions were resolved by consensus and are shown beneath the panel. (C) Capture Hi-C data is presented 5 Mbp across the HPV16 integration locus. The black line above the read peaks indicates the genomic window seen in panel B. Peaks of reads indicate regions of the host interacting with the integrated virus in three-dimensions. Long-range interactions between the HPV16 genome and host regions were resolved by consensus and are shown beneath the panel. In each panel, the scale bar represents the normalised read count. Additionally, protein-coding genes are shown in the first track with the direction of each gene indicated by colour (red, forward; blue, reverse), followed by the alignment of ChIP-seq data from the NHEK cell line (ENCODE). Post-translational histone modifications of active chromatin (H3K27ac, H3K4me1, H3K4me2, H3K4me3; green), repressive H3K27me3 (red), DNaseI hypersensitivity sites (blue) and CTCF sites (purple) are shown. Coordinates presented for each window are indicated at the top of each figure.

**Figure 3.**
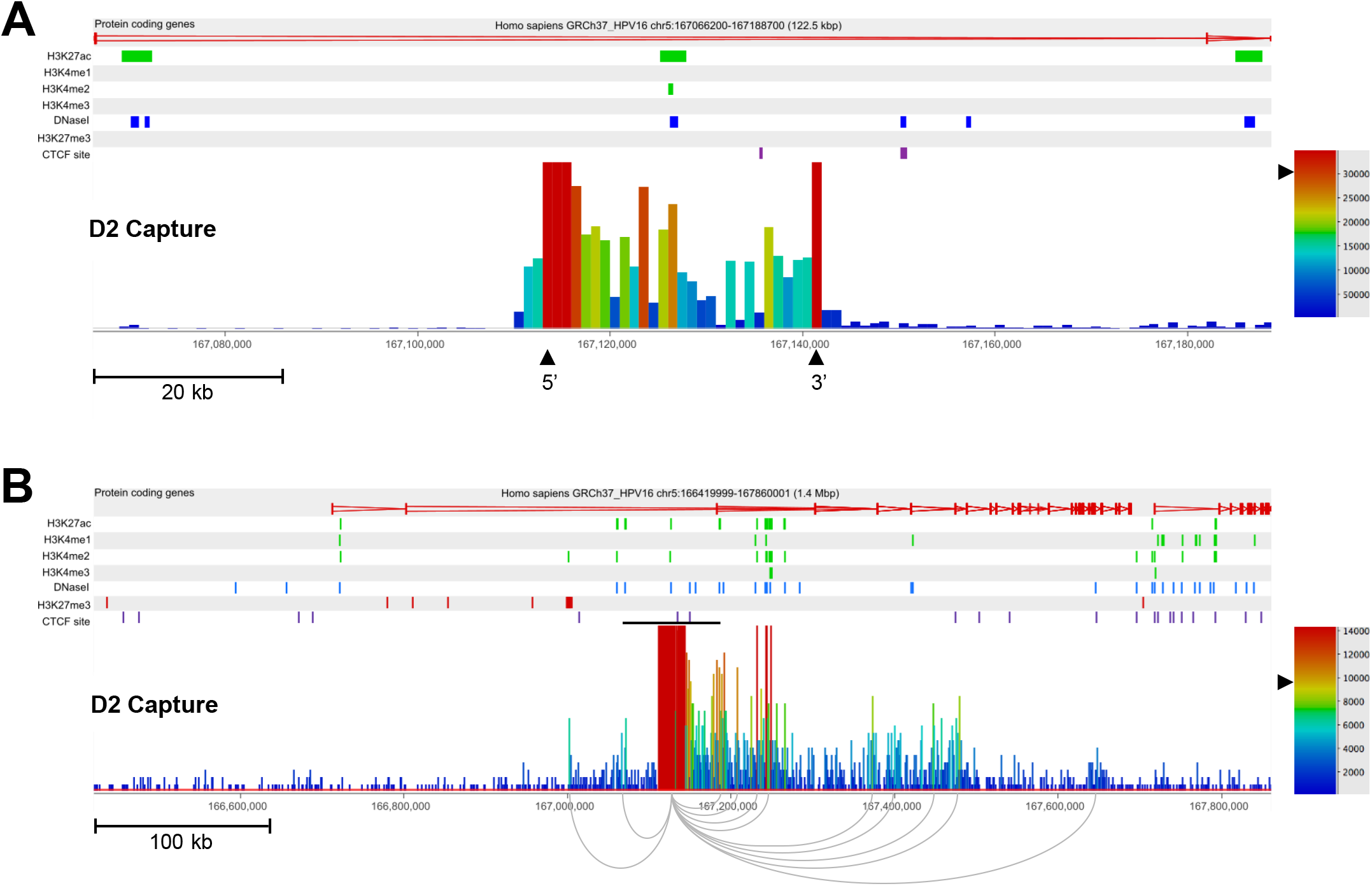
Identification of short- and long-range interactions between integrated HPV16 genomes and the host chromosome in W12 clone D2. (A) Capture Hi-C data is presented 122.5 kbp across the HPV16 integration locus. The 5’ and 3’ breakpoints of the virus are indicated by the tallest red bars and are labelled with black arrowheads. (B) Capture Hi-C data is presented 1.4 Mbp across the HPV16 integration locus. The black line above the read peaks indicates the genomic window seen in panel A. Peaks of reads indicate regions of the host interacting with the integrated virus in three-dimensions. Short-range interactions between the HPV16 genome and host regions were resolved by consensus and are shown beneath the panel. In each panel, the scale bar represents the normalised read count. Additionally, protein-coding genes are shown in the first track with the direction of each gene indicated by colour (red, forward; blue, reverse), followed by the alignment of ChIP-seq data from the NHEK cell line (ENCODE). Post-translational histone modifications of active chromatin (H3K27ac, H3K4me1, H3K4me2, H3K4me3; green), repressive H3K27me3 (red), DNaseI hypersensitivity sites (blue) and CTCF sites (purple) are shown. Coordinates presented for each window are indicated at the top of each figure.

Upon increasing the window range across the integration loci to ~700 kbp (with re-normalised read depth), peaks of 3D interaction outside of the initial 100 kbp window could be seen in clone G2 (Figure 2B) and clone D2 (Figure 3B). Multiple short-range (<500 kbp) 3D interactions were present ranging up to ~240 kbp and ~360 kbp away from the site of integration in clones G2 and D2, respectively (Figure 2B & 3B). Interestingly, sites of high intensity of 3D interaction with the host genome in both cell lines (Figure 2B & 3B, red bars) overlapped with histone marks of transcriptional activation or enhancers and DNaseI hypersensitivity sites, whereas these sites only correlated with host CTCF binding sites in clone G2 (Figure 2B, purple dashes). The presence of a long-range interaction (>500 kbp) was seen at this scale in clone D2 ~530 kbp from the integration site with the 3’ end of host gene *TENM2* (Figure 3B). However, long-range interactions in clone G2 were only visible when expanding the window range to 5Mb, whereupon two clear 3D interactions were determined downstream of the integration site with the furthest ~900 kbp from the HPV16 integration site (Figure 2C). This interaction was located at Chr5:53,520,000 within the first intron of host gene *ARL15,* coinciding with a cluster of host CTCF binding sites (Figure 2C, purple dashes).

To validate that direct interaction occurred between the integrated HPV16 genome and *ARL15* intron in clone G2, 3D DNA fluorescent in situ hybridisation (FISH) was carried out (Figure 4). Three fluorescent DNA probes were produced to hybridise to either the integrated HPV16 genome, the *ARL15* site of interaction, or a control region of the host genome with the same linear distance in the opposite direction (Figure 4A). Only cells containing one HPV16 signal and two copies of both the control and *ARL15* probes were analysed. A representative image from the resulting dataset (30,000 cells) is shown in Figure 4B and analysis of the 3D distances (x, y and z planes) indicated that, on the integrated chromosome, the HPV16 and *ARL15*-specific probes were significantly closer together than the HPV16 and control-specific probes (Figure 4C & D). Additionally, it was found that the distance between the control and *ARL15*-specific probes on the integrated allele was significantly lower than that on the unintegrated allele (Figure 4E & F), corroborating the finding that direct 3D interaction between the integrated HPV16 genome and *ARL15* gene does occur in clone G2.

**Figure 4.**
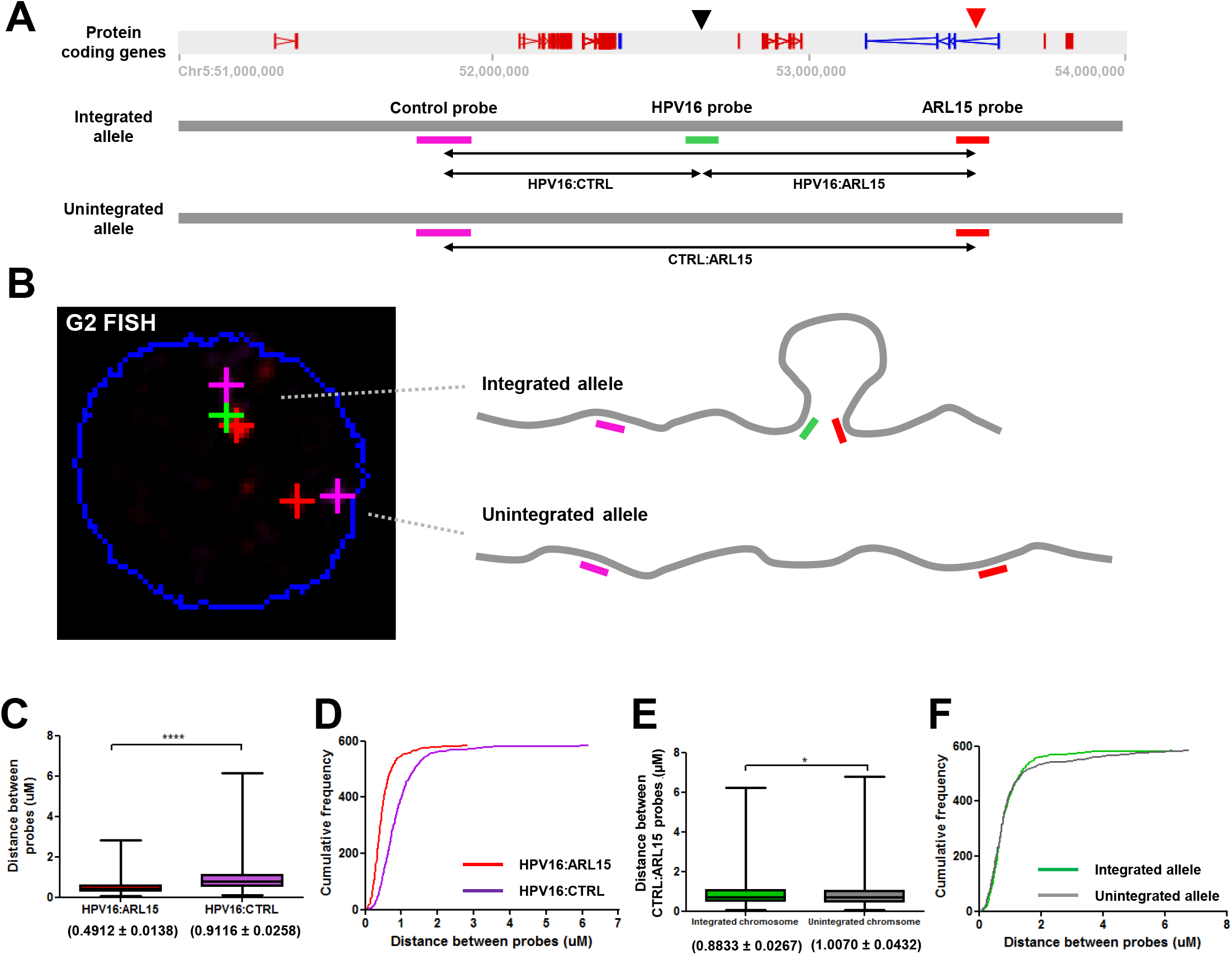
Validation of HPV16-host three-dimensional chromatin interactions in W12 clone G2 by FISH. (A) Schematic detailing the complementarity of the DNA probes used on the integrated and unintegrated alleles of a portion of chromosome 5 (51-54 Mbp) in W12 clone G2 to confirm interaction between the HPV16 genome (black arrow) and *ARL15* gene (red arrow): Control probe (51,676,020-51,873,551; purple), HPV16 probe (green), and ARL15 probe (53,473,886-53,584,235; red). Possible interactions between probe regions are also highlighted. (B) Representative image of the probes hybridised to W12 clone G2 genome of one cell in a 3D FISH experiment (nucleus boundary, blue) and interpretation of the associated chromosome spatial conformations. (C) Box-whiskers plot and (D) frequency distribution chart of the distance between both sets of FISH probes in the integrated allele of chromosome 5: HPV16:ARL15 (red box) and HPV16:control (purple box) (Mean ± SEM). (E) Box-whiskers plot and (F) frequency distribution chart of the distance between the Control and ARL15 probes in both the integrated (green) and unintegrated (grey) alleles (Mean ± SEM). n=585; * p<0.05, **** p<0.0001.

### HPV16 genome integration does not affect local host genome architecture

Since 3D interactions between integrated HPV16 and host genomes had been confirmed, we next asked whether these interactions could lead to structural changes in local nuclear architecture. Hi-C libraries, containing global host:host interactions, from both clones G2 and D2 were sequenced and, due to the different sites of integration on chromosome 5, these datasets were compared against each other within 5Mb windows spanning the two integration sites (Figure 5). Heatmaps of clone G2 Hi-C data (Figure 5A) showed an interaction profile consistent with data from clone D2 (Figure 5B), with interactions occurring across the 5Mb window including clearly defined regions of interacting host DNA up to ~1Mb, approximating to the size of a topologically associating domain (TAD). Insulation score analysis also determined no significant change across the 5Mb window between the two interaction profiles despite the presence of HPV16 genomes at the site in clone G2 (Figure 5C). Indeed, this finding was repeated when 2.5Mb either side of the clone D2 HPV16 integration site was compared to that of clone G2 (Figure 5D-F). Hence, HPV16 genome integration and interactions with the host chromosomes did not appear to be affecting the local nuclear architecture.

**Figure 5.**
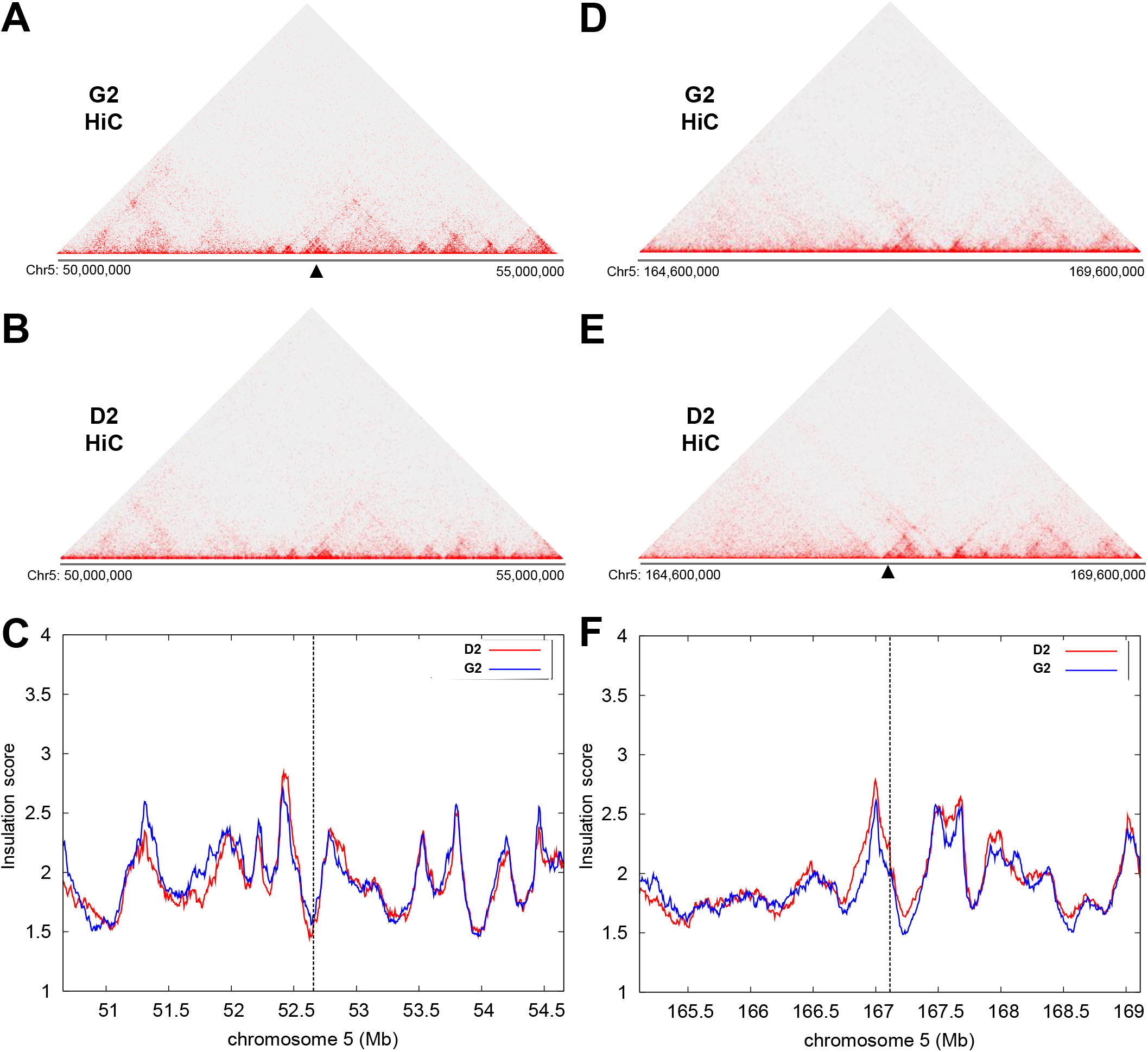
HPV16 genome integration does not disrupt host nuclear architecture at the integration site in W12 clones. (A, D) Hi-C data for W12 clone G2 is compared to (B, E) HiC data for W12 clone D2 using (C, F) an insulation score interpreting the association of topologically associating domains (TADs) for HPV16 integration sites within W12 clone G2 (Chr5: 50-55 Mbp; left column) and W12 clone D2 (Chr5: 164.6-169.6 Mbp, right column) showing no significant change to either window. Black arrow = integration site.

### HPV16 genome integration does modulate local host gene expression

As HPV16 genome integration, and resulting 3D interactions with the host genome, did not appear to affect local nuclear architecture, we sought to determine how far within the host:host chromosome interactions ‘loops’ from the HPV16 genome occurred. To the existing Region Capture Hi-C datasets, we aligned publically available TAD boundaries determined from two human cell lines(48) and found that all HPV16:host interactions occurred within the TAD of integration (Figure 6). Analysis of clone D2 found that the majority of HPV16:host genome interactions occurred within the TAD of integration, with only the long-range loop occurring outside of this range in an adjacent TAD (Figure 6C), although there is a possibility that all loops may exist here within the TAD of integration given differential calling of TAD boundaries across cell lines as all loops occur exclusively with the *TENM2* gene.

**Figure 6.**
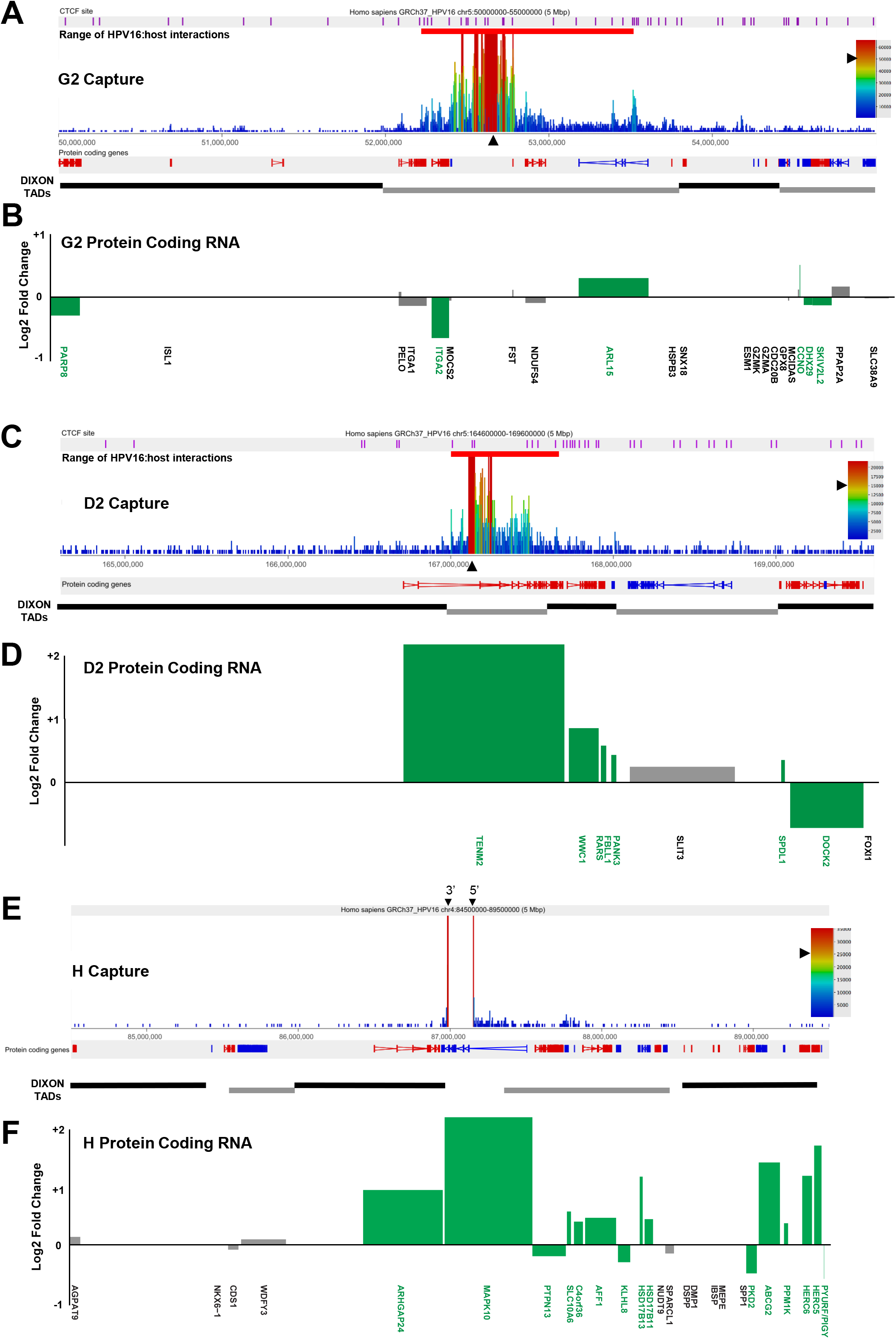
HPV16 genome integration leads to significant modulation of host gene expression in W12 clones G2, D2 and H. Capture Hi-C data is presented across (A-B) Chr5:50-55 Mbp for clone G2, (C-D) Chr5:164.4-169.6 Mbp for clone D2 and (E-F) Chr4:84.5-89.5 Mbp for clone H. HPV16 integration site is indicated with a black arrow and CTCF sites (purple) aligned across the top of the panel. Aligned protein coding genes are shown in the top track (rightward, red; leftward, blue) with the extent of topologically associating domains (TADs) determined by Dixon et al. shown below. (B, D, F) Charts indicating the transcript level of host protein coding genes within the 5 Mb region of integrant clones relative to a mean control level across all other clones. All data is shown as a Log2 fold change with significant changes indicated by green bars. Gene length is indicated by width of the corresponding bar.

Next, we addressed whether HPV16 genome integration and interactions with the host chromosome led to any changes in local host gene expression. Transcribed RNAs across a 5Mb window spanning the HPV16 integration site were assessed through comparison of total transcriptome RNA-seq data from an individual clone to an average of all other available datasets from W12 integrant clones with a different integration site. Analysis of protein coding RNAs from clone G2 found that host genes were both up- and down-regulated across the integration locus and in some cases unchanged, with no restriction to TADs (Figure 6B); a consistent finding across all cloned cell lines analysed. Despite all changes being less than ±2-fold change here, some genes were found to be significantly down-regulated (*PARP8, ITGA2, CCNO, DHX29, SKIV2L2*) while interestingly the only significantly up-regulated gene was that with a confirmed HPV16:host interaction, *ARL15* (1.23-fold; *p*< 0.05) (Figure 6B, green labelled genes). Compellingly, upon comparative analysis of Hi-C libraries between clones G2 and D2, a decrease in a host:host interaction was found within the TAD of integration (Supplementary Figure 12A, blue triangle), aligned with the HPV16:host interaction with the *ARL15* gene. Hence, changes of gene expression within TADs are possibly due to modulated host:host interactions at this level.

Analysis of host gene expression at the HPV16 integration site of clone D2 found again that transcription was both up- and down-regulated, while some genes appeared unaffected, with all but one modulation statistically significant (Figure 6D). The host gene into which HPV16 had integrated and exclusively interacted with, *TENM2*, was up-regulated 4.79-fold in comparison to the control average. No significant changes in host:host interaction were found here within the TAD of HPV16 integration or the adjacent TADs (Supplementary Figure 13). Interestingly, all other clones that exhibited HPV16 integration within a host gene showed up-regulation of expression of that gene. Expression of *RASSF6* in clones F and A5 increased 1.64- and 1.62-fold, respectively (Figure 7B & D, respectively), with *p*<0.0001 in both cases. In clone H, despite deletion of some of the host coding exons, expression of *MAPK10* was increased 4.47-fold (Figure 6F), and chimeric HPV16:host RNA-seq reads showed that both breakpoint fusion transcripts and spliced transcripts from the integrated HPV16 genome into an adjacent host exon (ENSE00001811960; Chr4:86,952,584), including differential HPV16 exon expression, was the likely cause of the overall increase in expression levels of this gene (Supplementary Figure 14). Indeed, splicing events from the integrated HPV16 genome into the host was seen across three of the five integrant clones analysed (Supplementary Table 1), including clone G2 where intergenic HPV16 integration led to spliced fusion transcripts, verified through PCR and Sanger sequencing, with non-coding host DNA, presumably through cryptic splice acceptor sites (Supplementary Figure 15). Interestingly, all splicing events occurred with host DNA within the region of host DNA amplification flanking the HPV16 integration site in clone G2.

**Figure 7.**
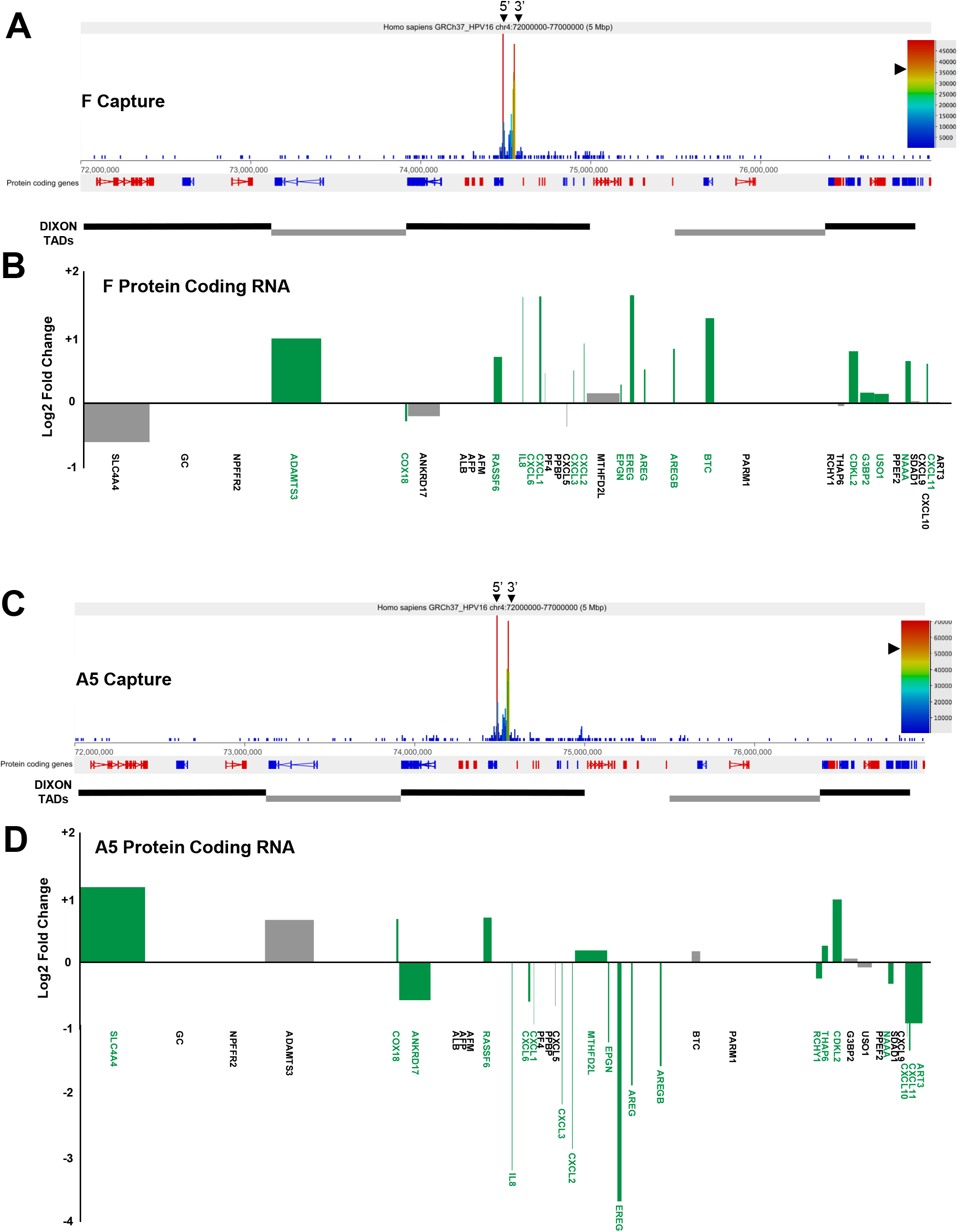
HPV16 genome integration leads to significant, but differential, modulation of host gene expression in W12 clones F and A5. Capture Hi-C data is presented across W12 clones (A-B) F and (C-D) A5 integration loci (Chr4: 72-77 Mbp). 5’ and 3’ ends of HPV16 integration site are indicated with black arrows. Aligned protein coding genes (rightward, red; leftward, blue) and the extent of topologically associating domains (TADs) determined by Dixon et al. are shown below. Charts indicating the transcript level of host protein coding genes within the 5 Mb region of W12 clones relative to a mean control level across all other clones. All data is shown as a Log2 fold change with significant changes indicated by green bars. Gene length is indicated by width of the corresponding bar.

### HPV16 genome integration modulates host gene expression across the chromosome

To further investigate the effect of HPV16 integration on host gene expression, the variance in gene expression in the genomic regions adjacent to the HPV16 integration site was compared with that of the whole chromosome. Again, comparing RNA-seq data from an individual clone to an average of all other available datasets from W12 integrant clones with a different integration site, host transcription variance (from both protein coding and non-coding regions) was calculated by grouping five adjacent genes into ‘bins’ and then comparing the range and variance within each bin to that of a mean level from across the whole chromosome (Figure 8 & Supplementary Figure 16). Expression of host genes across all regions appeared highly variable, with clones G2 (Figure 8A), F and A5 (Supplementary Figure 16C & E) all having bins with ±2-fold change. Multiple bins at and adjacent to the HPV16 integration site were also found to have highly significant (*p*<0.05 and *p*< 0.001) gene expression variance (Figure 8B & D & Supplementary Figure 16B, D & F). Hence, integration of HPV16 genomes into host chromosomes appears to have modulatory effects on the host gene expression far beyond the immediate locus of integration.

**Figure 8.**
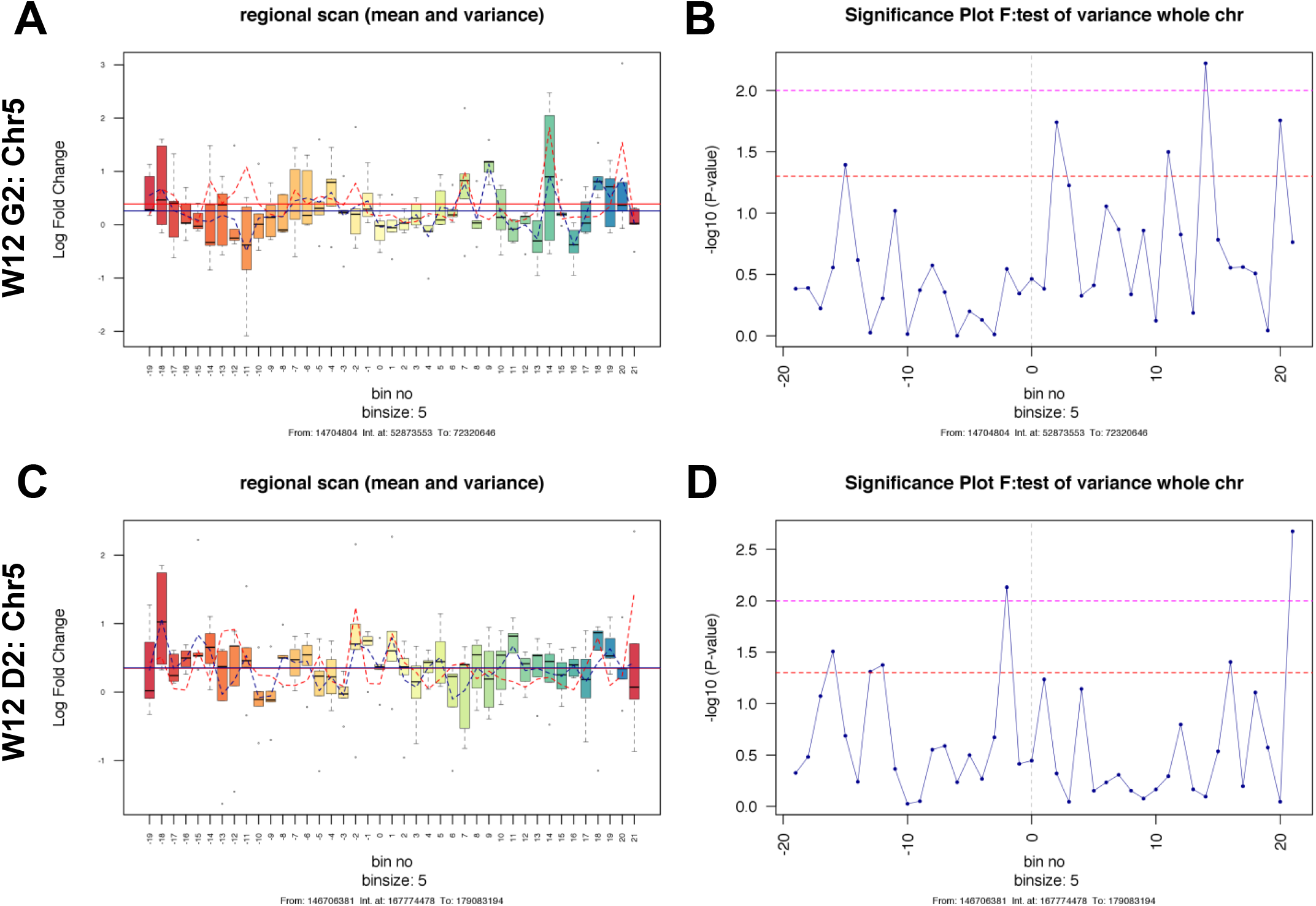
Variance in host gene expression across the host genomic region containing the HPV16 integration site in W12 clones G2 and D2. Each left panel indicates the range and variance of host gene expression in W12 integrant clones [A) W12 G2, and C) W12 D2], focussing on 100 genes either side of the HPV16 integration site. For each clone, gene expression levels were compared with a 6-clone integrant average control level. In each panel, the HPV16 integration site is centred on ‘bin 0’. Each bin contains five genes, with no overlap between bins. The box and whisker plots illustrate the range of gene expression levels within each bin, with the bar indicating median values, the box the IQR and the whiskers the range. The mean gene expression across the whole chromosome is indicated by the solid blue line, while the mean level of gene expression across individual bins is shown by the dotted blue line. The mean variance of gene expression across the whole chromosome is indicated by the solid red line, while the mean level of gene expression across individual bins is shown by a dotted red line. Each right hand panel shows the significance of the variance in gene expression within each bin [B) W12 G2, and D) W12 D2]. Each point represents a five-gene bin, corresponding to those in the lefthand panels. The horizontal lines indicate the significance of the variance in each bin, compared with the variance in gene expression across the whole chromosome (above the dashed red line, p<0.05; above the dashed pink line, p<0.01).

## Discussion

Progression of disease toward cervical carcinoma is markedly associated with integration of HPV genomes into host chromosomes whereby dysregulation of the control of HPV oncogene expression occurs. Our previous work has shown that, across five integrant cell lines cloned from the W12 model system (F, A5, D2, H & G2), levels of virus transcript per template correlate with multi-layered epigenetic changes that regulate transcription from the integrated HPV16 genomes(15, 21). However, it has not been determined how integration at these sites may produce a more or less selectable clone during outgrowth in our model, which mirrors the natural process of cervical carcinogenesis. To answer this, we developed a state-of-the-art technique ‘HPV integrated site capture’ (HISC) to determine the precise loci of HPV16 integration sites and utilised Region Capture Hi-C to determine if 3D interactions between HPV16 and host genomes could be involved in modulating host gene expression, thereby driving selection of certain clones. With this new technology, we first sought to confirm the integration sites of HPV16 genomes in our five clones through next generation sequencing (NGS) of samples.

In all the W12 integrant clones tested, the HPV16 genome was shown to interact with regions of host chromosomes in *cis*; there were no examples of HPV16 interacting with the host in *trans*, although this could be due to limited sequencing depth. These interactions occurred from across the HPV16 genome, although there was an absence of any interactions from a large proportion of the *E1* gene. This was due to technical reasons associated with the design of the RNA baits, based upon *MboI* restriction sites within the HPV16 genome, rather than a true biological finding. These interactions pinpointed the definitive loci of integration of the HPV16 genomes within each W12 integrant clone, with PCR and sequencing confirming that these locations differed from their original published sites(15, 21, 24) in all but one clone (clone H). More interestingly, we found that clones F and A5 had the same HPV16 integration site, with the same HPV16 breakpoints and inversion, likely indicating that one cell line is a precursor of the other since they exhibit very different phenotypical characteristics(15, 21, 24). Our data also support the growing understanding that integration of HPV genomes takes place, at least to some degree, through microhomology-mediated repair (MHMR) of DNA breaks(25, 26, 29, 32, 47, 49-51) as our precise determination of HPV16:host breakpoints through HISC, RNA-seq and Sanger sequencing allowed interrogation of the flanking sequences at each junction showing higher levels of homology between HPV16 and host sequences than expected. Indeed, this support for MHMR lends further support to the theory that HPV integration occurs via two main processes: ‘direct’ and ‘looping’ integration(32, 47). Here, deletion of a proportion of the host genome adjacent to the HPV16 integration site in W12 clone H is consistent with direct integration and, for the first time as far as these authors are aware, we have shown that deletion can occur in a homozygous fashion. All other clones examined (F, A5, D2, G2) showed clear signs of host genome amplification flanking the integration site that would be consistent with looping integration from which integrants are now sometimes known as ‘type III’ (22, 34). There remains the possibility that these regions could be amplified as extra-chromosomal virus-host fusion episomes maintained by the HRHPV origin of replication, which has been proposed following analysis of HNSCC TCGA datasets elsewhere(29, 52), although our techniques here appear to consistently illustrate canonical chromosomal integration.

Despite others’ previous analysis of integration sites primarily focussing on cervical SCCs(12), we have also shown in our W12 integrant cell lines, which reconstitute the early stages of cervical carcinogenesis, that HPV16 integration, although occurring in both intronic and intergenic regions in our population, appears more readily to occur within regions of increased chromatin accessibility (DNAseI hypersensitive regions) and with high association of transcriptional enhancers (H3K4me1/H3K27ac) or activity (H3K4me2/3), as found previously(12, 53-55). Intriguingly, an accomplished recent analogous study of one W12-derived subclone (20861), which has 26 tandemly integrated HPV16 genomes interspersed with 25 kb of flanking cellular DNA at chromosome 2p23.2 (a so-called type III integrant), has shown that establishment of super-enhancer like regions can occur through ‘looping’ integration resulting from amplification of both virus LCR and a basal cellular enhancer. This leads to an enrichment of super-enhancer marks H3K27ac and BRD4, and likely drives high expression of virus *E6/E7* fusion transcripts with subsequent selection and neoplasia(33, 34).

Highly specialised Region Capture Hi-C/HISC analysis of the W12 integrant clones here confirmed that 3D interactions, both short-range (<500 kbp) and long-range (>500 kbp), between integrated HPV16 genomes and host chromatin, inferred elsewhere previously(22, 38), do occur in cell lines that mirror the very early stages of cervical carcinogenesis. We were able to verify that these ‘loops’ are present in 3D using DNA FISH analysis in one of our cell lines, clone G2, by confirming the distances between the integrated HPV16 genome and the site of interaction in the *ARL15* gene in comparison to that of a control region. Interestingly, the sites of interaction on the host genome both in clone G2 and D2 more often than expected aligned with sites of interaction of the host architectural protein CTCF. It is known that the HPV genome is able to interact with CTCF through virus-specific sequences(56), contributing to differentiation-dependent control of virus gene expression through loops with YY1(57), and it is appealing to hypothesise that insertion of an ectopic CTCF-binding site into the host genome through HPV genome integration, as has been found with the human T-cell lymphotropic (HTLV-1)(58, 59), may lead to modulation of the local host genome architecture and changes in host gene expression. Indeed, even with the loss through integration of the HPV16 *E2* located CTCF-binding site in clone G2, interaction could still be driven by further putative sites in the *L2/L1* genes(56) or could occur through promoter/enhancer-like interactions from the HPV genome as is believed to be the situation in the HPV18 integrant HeLa cell line(38). This is supported by our finding that loops between the integrated HPV16 genome in our clones also appear to localize to host sites of transcriptional activity and chromatin marks of promoters/enhancers. Whether the orientation of the HPV genome after integration has any effect on directionality of these interactions or loops and indeed whether interaction with these host chromatin domains has any effect on transcript levels produced by the integrated HPV16 genome remains unknown.

Here, HPV16 integration into a gene (*MAPK10* in clone H; *TENM2* in clone D2; *RASSF6* in clones F and A5) clearly caused increases in expression of that host gene. Despite the loss of some coding exons in *MAPK10* in clone H, direct co-linear insertion of a HPV16 genome led to splicing from the HPV16 genome into the next host exon. Thus, although overall levels of MAPK10 transcript were raised, this was due to an increase in the production of RNA from 3’ end exons and possible fusion to HPV16 transcripts. It remains to be determined whether these truncated and/or fusion transcripts could code for protein or indeed whether any expressed protein would be functional. Regardless of integration mechanism, the presence of at least one integrated copy of an HPV16 genome caused modulation of host gene expression across a wide range of that chromosome. Statistical analysis of groups of host genes, including those at the site of integration, in comparison to other W12 integrant cell lines showed wide ranging influence on the host gene expression profiles with many sites having >±2-fold changes in transcript level.

Our analysis of total host:host 3D interactions through interrogation of Hi-C libraries did not appear to provide evidence that host topologically associating domains (TADs) were greatly affected by HPV16 genome integration. Interestingly, almost all 3D interactions between HPV16 genomes and the host chromosome occurred within a single TAD, leaving the hypothesis that TAD boundaries may in some way be able to inhibit HPV:host interactions into adjacent TADs. This barrier to more elongated interactions may also stretch to *trans* interactions with chromosomes other than that of integration, although this remains unverified. Regardless, looping from the integrated HPV16 genome in clone G2 to the *ARL15* gene was associated with an increase in the level of its transcript. Further comparative analysis of our Hi-C data sets showed that this interaction may in fact cause a decrease in the usual host:host interaction between *ARL15* and a region upstream of the HPV16 integration site. Interestingly, the host genes around this upstream interaction site were largely down-regulated. Therefore, it is possible that the relatively stronger interaction between the HPV16 promoter/enhancer and *ARL15* gene that might up-regulate ARL15 transcript levels may supersede usual host interactions, some of which may be promoter:enhancer interactions that maintain regular host transcript levels. This modulation of host gene expression distal from the HPV genome integration site is in line with the current theory of Viro-TADs and host gene expression changes(22, 59, 60).

Although we are not the first to use Capture technology to precisely determine HPV integration sites(22, 26, 34, 47, 50, 51, 61), our analysis of Region Capture Hi-C and full host:host Hi-C libraries have allowed the first definitive determination of 3D interactions between integrated HPV and host genomes with local (intra-TAD) changes to usual local host:host interactions. The downstream effect of these modulations to local host architecture is modulation of the host gene expression program, at least within the same TAD. Therefore, integration near to or within a ‘cancer-causing gene’ does not appear essential to influence such genes due to these *cis*-driven distal events. However, it remains to be fully determined how these HPV:host 3D interactions are initiated and maintained, and whether this type of interaction or the downstream effect on host gene expression is selected for, as would be hypothesised through evidence provided by HeLa cells(22, 38), as indeed whether these interactions also drive the level of HPV transcripts from the integrated virus genome.

## Materials and Methods

### Cell culture

Detailed descriptions of the W12 system have been published previously(10, 41, 62) including W12 integrant clone generation(15, 24). The five W12 clones used here were episome-free, did not express the HPV16 transcriptional regulator E2(15) and were grown in monolayer culture in order to restrict cell differentiation and maintain the phenotype of the basal epithelial cell layer(63). Additionally, W12 clones were analysed at the lowest possible passage after cloning (typically p3 to p8) in order to minimise any effects of genomic instability caused by deregulated HPV16 oncogene expression(21).

### HPV16-host breakpoint and splice junction confirmation

To verify chimeric DNA sequences of HPV16-host breakpoints determined by Capture-seq, and to confirm splice junction sequences from clone G2 RNA-seq analysis, primers were designed using either Primer3 (Primer3Web) alone or Primer-BLAST (NCBI) specific to the DNA sequence (or cDNA sequence from reverse transcribed clone G2 RNA samples (QuantiTect Reverse Transcription Kit, Qiagen)) for PCR (PCR SuperMix High Fidelity, Thermofisher). PCR products were gel extracted and then Sanger sequenced using both 5’- and 3’-end primers to confirm reads from each end of the product. Each analysis was carried out in duplicate. Primer pairs used for PCR of HPV16-host breakpoints and clone G2 splice junctions are given in Supplementary Table 2 and 3, respectively.

### qPCR of HPV16-host breakpoints and genomic DNA

Primers were designed using either Primer3 alone or Primer-BLAST (NCBI) specific to the chimeric DNA sequence of HPV16-host junctions determined by Capture-seq to verify integration sites, as well as host DNA spanning integration sites to determine copy number after HPV16 integration. Primers used for qPCR are given in Supplementary Tables 4 and 5. Host DNA copy number was quantified by comparison to TLR2 and IFNβ, as reported previously(15). Conditions used for all primer pairs on an Eppendorf Mastercycler Realplex were: 95°C for 2min; 40 cycles of 95°C for 15sec, 58°C for 20sec, 72°C for 15sec, 76°C for 5sec and read; followed by melting cure analysis from 65°C to 90°C to confirm product specific amplification.

### Insulation score plots

Measurement of the topological domain structure along the chromosomes was computed with an average insulation score profile at the TAD boundaries. The insulation score is the standardized -log enrichment of contacts between the downstream and upstream 300kb regions (-log (a/(a+b1+b2)) where a is the number of contacts between, and b1 and b2 the number of contacts within the upstream and downstream 300kb regions). Using this definition, a more positive insulation score indicates a stronger TAD boundary.

### DNA Fluorescence In-Situ Hybridisation (FISH)

BAC clones RP11-467N14 (control locus) and CTD-2015C9 (ARL15 locus) were purchased (Thermofisher), whereas the HPV plasmid pSP64-HPV16 was prepared in house. BAC/plasmid DNA was purified using the NucleoBond BAC100 kit (Macherey-Nagel), and labelled with aminoallyl-dUTP by nick translation. After purification, 0.5-1 μg labelled BAC DNA was coupled with Alexa Fluor 488, Alexa Fluor 555 or Alexa Fluor 647 reactive dyes (Life Technologies) according to the manufacturer’s instructions, and DNA FISH was performed as described elsewhere(64).

### Chromatin crosslinking

Formaldehyde crosslinking of 30 million cells was performed by supplementing standard EGF positive culture medium with formaldehyde to a final concentration of 2% and was carried out for 10 min at room temperature. Crosslinking was quenched by the addition of ice-cold glycine to a final concentration of 125 mM. The adherent cells were scraped from the cell culture plates after crosslinking, collected by centrifugation (400 g for 10 minutes at 4°C), and washed once with PBS (50 ml). After centrifugation (400 g for 10 minutes at 4°C), the supernatant was removed, and the cell pellets were snap-frozen in liquid nitrogen and stored at −80°C.

### Hi-C library generation

Cells were thawed on ice, and then lysed on ice for 30 minutes in 50 ml freshly prepared ice-cold lysis buffer (10 mM Tris-HCl pH 8, 10 mM NaCl, 0.2% Igepal CA-630, one protease inhibitor cocktail tablet (Roche complete, EDTA-free)). Following the lysis, nuclei were pelleted (650 g for 5 minutes at 4°C), washed once with 1.25 x NEBuffer 2, and then resuspended in 1.25 x NEBuffer 2 to make aliquots of 5-6 million cells for digestion. SDS was added (0.3% final concentration) and the nuclei were incubated at 37°C for one hour (950 rpm). Triton X-100 was added to a final concentration of 1.7 % and the nuclei were incubated at 37°C for one hour (950 rpm). Restriction digest was performed overnight at 37°C (950 rpm) using 800 units MboI (NEB) per 5 million cells. Restriction fragment ends were filled in using Klenow (NEB) with dCTP, dGTP, dTTP and biotin-14-dATP, and the blunt-ended DNA was ligated following the in-nucleus ligation protocol described previously (Nagano et al., 2015), with minor modifications. Prior to ligation, excess salts and enzymes were removed by centrifugation (600 g for 5 minutes at 4°C) and the cell pellet was re-suspended in 995 μl of 1 x ligation buffer (NEB) supplemented with BSA (100 μg/mL final concentration). The ligation was carried out using 2000 units of T4 DNA ligase (NEB) per 5 Mio starting material of cells, at 16°C for 4 hours, followed by 30 min at room temperature. Chromatin was then de-crosslinked overnight at 65°C in the presence of proteinase K (Roche), purified by phenol and phenol-chloroform extractions, precipitated with ethanol and sodium acetate and re-suspended in TLE (10 mM Tris-HCl pH 8.0; 0.1 mM EDTA). The DNA concentration was measured using the Quant-iT PicoGreen assay (Life Technologies). 40 μg of Hi-C library DNA were incubated with T4 DNA polymerase (NEB) for 4 hours at 20°C to remove of biotin from non-ligated fragment ends, followed by phenol/chloroform purification and DNA precipitation overnight at −20°C. DNA was sheared to an average size of 400 bp using the Covaris E220 (settings: duty factor: 10%; peak incident power: 140W; cycles per burst: 200; time: 55 seconds). End-repairing of the sheared DNA (using T4 DNA polymerase, T4 DNA polynucleotide kinase, Klenow (all NEB)) was followed by dATP addition (Klenow exo-, NEB) and a double-sided size selection using AMPure XP beads (Beckman Coulter) to isolate DNA ranging from 250 to 550 bp. Biotin-marked ligation junctions were immobilised using MyOne Streptavidin C1 Dynabeads (Invitrogen) in binding buffer (5 mM Tris-HCl pH 8.0, 0.5 mM EDTA, 1M NaCl) and after stringent washing in the same buffer at 55°C for 10 min ligated to the custom SCRiBL adapter using 1600 units of T4 DNA ligase (NEB) for 2 hours at room temperature. These adapters were generated by annealing SCRiBL_adapter_1 and SCRiBL_adapter_2 (table X). The immobilised Hi-C libraries were amplified using the custom primers PE PCR 1.0.33 and PE PCR 2.0.33 with 7-9 cycles. After PCR amplification, the Hi-C libraries were purified with AMPure XP beads (Beckman Coulter). Quantity and integrity of the Hi-C libraries was determined by Bioanalyzer profiles (Agilent Technologies).

### Genomic DNA library generation

Cells were thawed, lysed and nuclei were isolated as described above. Nuclei from 5-6 Mio cells were treated with SDS and Triton X-100 as described for the generation of Hi-C libraries. All Hi-C specific steps, such as MboI digestion, restriction fragment end fill-in, blunt end ligation and the removal of biotin from un-ligated restriction fragment ends were mock performed by replacing the respective enzymes with an equal amount of water. All other steps were performed as described for the generation of the Hi-C libraries. The biotin-streptavidin pull down was omitted and, therefore, the ligation of the custom sequence adapters was done in solution by adding 4 μl adaptors (30 μM) and 1600 units T4 DNA ligase (NEB). The ligation was carried at for 2 hours at room temperature on a rotating wheel in 1x ligation buffer (NEB). Pre-capture PCR amplification was carried out using the custom primers PE PCR 1.0.33 and PE PCR 2.0.33 with 7-8 cycles. The amplified libraries were purified with AMPure XP beads (Beckman Coulter) and the quantity and the quality was assessed by Bioanalyzer profiles (Agilent Technologies).

### Capture RNA bait library design

120-mer capture RNA baits were bioinformatically designed to both ends of MboI restriction fragments overlapping the HPV16 genome. Requirements for target sequences were as follows: GC content between 25% and 65%, no more than two consecutive Ns within the target sequences, and maximum distance to a MboI restriction site 330 bp. For short MboI fragments, where 120-mer RNA baits originating from both ends would have overlapped (potentially interfering with optimal hybridization to Hi-C libraries), only the Watson (coding or sense) strand was used for capture RNA bait design, and if necessary the baits were trimmed to minimum length no shorter than 97 nt. This resulted in the design of 16 RNA bait sequences (Supplementary Table 6) covering the MboI restriction fragment ends of the entire HPV16 genome, with the exception of two fragments too short (18 and 63 bp, respectively) for capture RNA bait design.

### Biotinylated RNA bait library for Region Capture Hi-C generation

The process of HPV16-specific Region Capture Hi-C was carried out essentially as previously published(65). DNA sequences encoding for the 16 RNA bait sequences, with different restriction enzymes sites on each side (BglII on one site and either HindII or SpeI on the other), which were separated by a 3 bp random spacing sequence, were ordered as two gBlocks® (Integrated DNA Technologies, supplementary table X) and cloned into plasmid vectors using the Zero Blunt® TOPO® cloning kit with One Shot® TOP10 Chemically competent cells according to manufacturer’s instructions. Both gBlocks were extracted from plasmid DNA by EcoRI (30 units) restriction enzyme digestion at 37°C for 2 hours. Having a BglII and another restriction enzyme sites on the other end, enabled BglII side specific ligation of a T7 promoter sequence adapter with a BamHI overhang essentially as per manufacturer’s instructions, preventing the generation of overlapping complementary transcripts. These adapters were generated by annealing T7_promoter_adapter_1 and T7_promoter_adapter_2 as per manufacturer’s insctructions. Digestion with both restriction enzymes and adaptor ligation were done in one reaction simultaneously, in the presence of BamHI. Two reactions containing 700 ng of gBlock1 DNA or 850 ng of gBlock2 DNA, 30 units BglIl each, 100 units BamHI each, 5-fold molar access of pre-annealed T7 promoter adapters and either 80 units HindIII (NEB) or 40 units SpeI (NEB) were incubated at 37 °C for 2 hours in 1x T4 DNA ligase buffer (NEB). Following this incubation 1200 units T4 DNA ligase (NEB) were added to each reaction and incubated at 25 °C for 3 hours. The samples were then run on a 1% agarose gel and specific bands at 180 bp were cut out and gel purified. Equimolar amounts of sequences were *in vitro* transcribed according to manufacturer’s instructions using the T7 MegaScript kit (Ambion) with biotin-labelled UTP (Roche). The RNA was then purified using the MEGAclear kit (Ambion) following the manufacturer’s instructions.

### Biotinylated RNA bait library generation for HISC

Four consecutive and non-overlapping fragments from the pSP64 HPV16 bacterial artificial chromosome were PCR amplified using the Expand High Fidelity PCR system (Roche). This resulted in complete coverage of the HPV16 genome. T7 promoter sequences (Roche) were added to one side of the PCR product during this PCR amplification, enabling subsequent directional *in vitro* transcription. Sequences were *in vitro* transcribed in the presence of biotin-UTP and purified as described above, followed by fragmentation to 120 nt with 4 nM magnesium chloride at 95°C for 7 min. Fragmented biotinylated RNA was purified by Isopropanol precipitation followed by two subsequent washes with 75% ethanol.

### Solution hybridization Region Capture of Hi-C libraries

500 ng to 2000 ng of Hi-C library DNA or genomic DNA library were concentrated using a vacuum concentrator (Savant SPD 2010, Thermo Scientific) and then re-suspended in 5 μl dH_2_O. 2.5 μg mouse cot-1 DNA (Invitrogen) and 2.5 μg sheared salmon sperm DNA (Ambion) were added as blocking agents. To prevent concatemer formation during hybridization, 1.5 μl blocking mix (300 μM) was added (equimolar mix of P5_b1_for_33, P5_b1_rev_33, P7_b2_for and P7B2_rev (see list X for sequences)). Biotinylated RNA baits were used in a ratio 1:12 to Hi-C libraries (25 ng biotinylated RNA baits per 300 ng of Hi-C library) and supplemented with 30 units SUPERase-In (Ambion). Biotinylated RNA baits for capture DNA-Seq were used in a ratio of 1:3.33 (300 ng RNA baits per 1,000 ng genomic DNA library) and supplemented with 30 units SUPERase-In. The DNA was denatured at 95°C for 5 min in a PCR machine (PTC-200, MJ Research; PCR strip tubes (Agilent 410022)) and then incubated with the biotin capture RNA at 65°C, in hybridization buffer (5 x SSPE (Gibco), 5 x Denhardt’s solution (Invitrogen), 5 mM EDTA (Gibco), 0.1 % SDS (Promega)) for 24 hours, in a total reaction volume of 30 μl. Captured DNA/RNA hybrids were enriched using Dynabeads MyOne Streptavidin T1 beads (Life Technologies) in binding buffer (1 M NaCl, 10 mM Tris-HCl pH 7.5, 1 mM EDTA) for 30 minutes at room temperature. After washing (once in wash buffer 1 (1 x SSC, 0.1 % SDS) for 15 minutes at room temperature, followed by three washes in wash buffer 2 (0.1 x SSC, 0.1 % SDS) for 10 minutes each at 65°C), the streptavidin beads (with bound captured DNA/RNA) were re-suspended in 30 μl 1 x NEBuffer 2. Post-capture PCR amplification was carried out using between six to nine cycles using primer pairs that consisted of one TruSeq adapter reverse compliment and the TruSeq universal adapter (see table X for sequences) from streptavidin beads in multiple parallel reactions, which were then pooled to purify the PCR products using AMPure XP beads (Beckman Coulter).

### Paired-end next generation sequencing

Two biological replicate Hi-C and capture Hi-C libraries were prepared for each of the cell lines. Sequencing was performed on Illumina HiSeq 2500 generating 50 bp paired-end reads (Sequencing Facility, Babraham Institute). CASAVA software (v1.8.2, Illumina) was used to make base calls and reads failing Illumina filters were removed before further analysis. Output FASTQ sequences were mapped to the human reference genome (GRCh37/hg19) containing the HPV16 genome as an extra Chromosome and were filtered to remove experimental artefacts using the Hi-C User Pipeline(66).

### Sequence analysis

#### HiCUP & SeqMonk

Sequence data was obtained from Illumina HiSeq paired-end sequencing. Using the HiCUP Pipeline(66) paired-end Capture Hi-C (cHi-C) fastq files were mapped with Bowtie 2(67) to a human GRCh37 reference containing a HPV16 pseudo-chromosome. HiCUP removes invalid and artefactual di-tags by overlaying the di-tags on an *in silico* restriction digest of the reference. The resulting BAM files contained putative di-tags for use in subsequent analyses. SeqMonk (https://www.bioinformatics.babraham.ac.uk/projects/seqmonk/) was used to quantitate and visualise the density of di-tags contained in the BAM files. The HPV16 sequence and annotation files were downloaded from the European Nucleotide Archive (www.ebi.ac.uk/ena/data). ENCODE Annotation for NHEK(68) was obtained from Ensembl release 75(69).

#### Circos

The raw cHi-C fastq files were converted to fasta format and BLAST(70) was used to search the HPV16 genome for reads mapping to it. The partner human reads were determined and the 2 sets of reads were mapped to the GRCh37 reference containing the HPV16 pseudo-chromosome using Bowtie 2. The BAM outputs were converted to BED format and modified to be compatible with the circular visualisation tool Circos(71). The HPV16 genome was split into bins of 500 bp and the count per bin determined from the chimaeric human-HPV16 ditags. The counts, the HPV16 MboI restriction map and gene coordinates were annotated on the Circos plots.

#### GOTHiC

The HiCUP output was converted to format compatible with the Bioconductor package GOTHiC(72). To find significant interactions between distal locations GOTHiC implements a cumulative binomial test based on read depth. This was used to identify regions of the human genome in contact with the HPV16 pseudo-chromosome at a resolution of 1kb. Di-tag mappings were visualised with Circos after filtering the previous Circos input by the GOTHiC determined interactions.

#### Breakpoint Mapping with USearch

The precise sites of HPV16 integration in the W12 cell lines were identified by sequencing HISC libraries. The raw fastq files were converted to fasta format and BLAST was used to search for reads mapping to the HPV16 genome. From these, the corresponding human tags were determined. Fast clustering of the reads with USearch(73), based on an sequence identity score of 0.65, identified clusters of sequences in the human and HPV16 derived reads. Consensus sequences from non-singleton clusters were obtained by aligning the clustered reads to each other using Clustal Omega(74). The breakpoints were inferred from these consensus sequences and validated by Sanger Sequencing(75). From the validated integration sites, custom chimaeric references were generated for each W12 line. Due to the existence of tandem amplifications in some of the regions of integration, two versions of the chimaeric human-HPV16 chromosomes were generated. In the first case, the HPV16 provirus was 5’ of a single amplified human sequence. For the second, the provirus was placed 3’ of the amplified human sequence. For another W12 line, ‘H’, there is a deletion in the region of integration and this was reflected in the chimaeric chromosome.

#### Juicer and Juicebox

Using the specific chimaeric references, Hi-C contact maps at different resolutions were generated from raw Hi-C fastq files using the Juicer Pipeline(76). Juicer constructs a compressed contact matrix from pairs of genomic positions located in close proximity in 3D space. The Hi-C contact maps were imported into Juicebox(77) for visualisation.

#### RNA-seq analysis and alignment

Complementary DNA (cDNA) libraries were prepared for the five W12 clones (two biological replicates each) by total RNA extraction from confluent cells with Ribo-Zero rRNA depletion and DNAse treatment before cDNA was prepared with the TruSeq RNA and DNA Sample PrepKit (Illumina). 50bp paired-end cDNA libraries sequenced on an Illumina HiSeq 2000 (Genomics Core Facility, EMBL Heidelberg). Sequence adapters were trimmed from the reads with Kraken(78). Trimmed FASTQs were mapped against a GRCh37.p13 reference transcriptome (Ensembl version75) that included HPV16 transcript annotation using STAR(79) with default parameters. Strand specific gene counts were obtained from alignments with HTSeq(80) and differential gene expression analysis performed using the R/Bioconductor package DESeq2(81). Modulation of host transcript levels due to virus integration was then evaluated per clone in comparison to the mean expression of all other clones.

## Data availability

All data supporting the findings of this study are present within the article and its Supplementary Information files, with all sequencing data deposited in the ArrayExpress database at EMBL-EBI (www.ebi.ac.uk/arrayexpress)(82) under accession numbers: E-MTAB-10152; E-MTAB-10154; E-MTAB-10155.

## Acknowledgements

Research was funded by a Cancer Research UK Programme Award (A13080). ELAD was supported by a PhD studentship from The Pathological Society of GB & NI awarded to IJG and NC.

## Author contributions

IJG and NC conceived the study. IJG, ELAD, MM, SS, CGS and NC devised the experimental approach. ELAD, MM and IJG performed the experiments. ELAD, IJG, MM, JMM, GB, SPS, CGS, SS, CV, PF, AE and NC analysed the data. IJG, ELAD and NC wrote the manuscript.

