## Supplementary figures and legends for "Three-dimensional interactions between integrated HPV genomes and cellular chromatin dysregulate host gene expression in early cervical carcinogenesis"

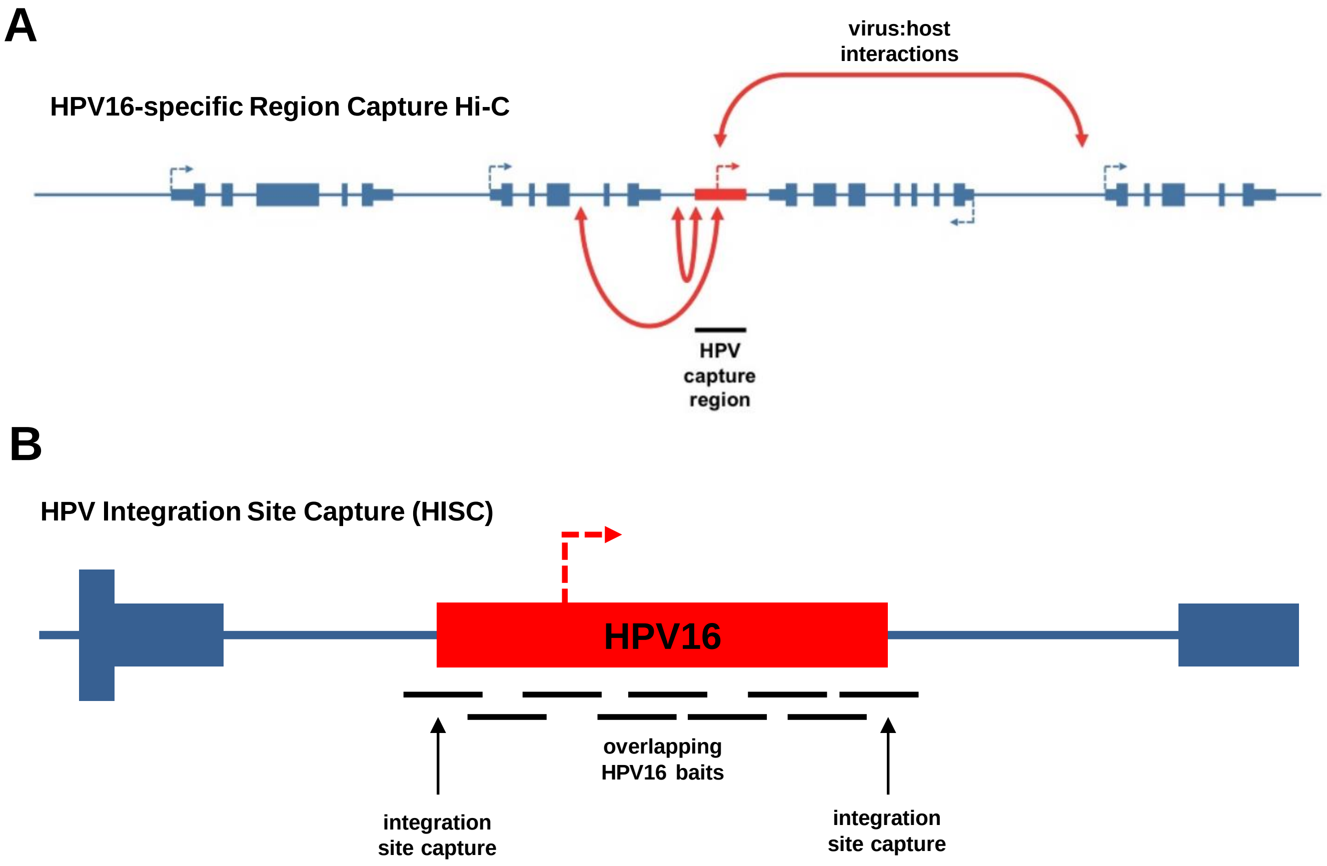


**Supplementary Figure 1. Diagram illustrating the principles of 'HPV16-specific Region Capture Hi-C' and 'HPV Integration Site Capture' (HISC) .** (A) HPV16-specific baits (consolidated pictorially as black line) are used to isolate both short- and long-range 3D interactions (red double headed arrow) between a capture region (integrated HPV16 genome) and the host genome. (B) HPV16-specific baits (black lines) are used to enrich HVP16:host breakpoints from an 'undigested' sequencing library. (blue indicates host chromosome, genes and promoters.)

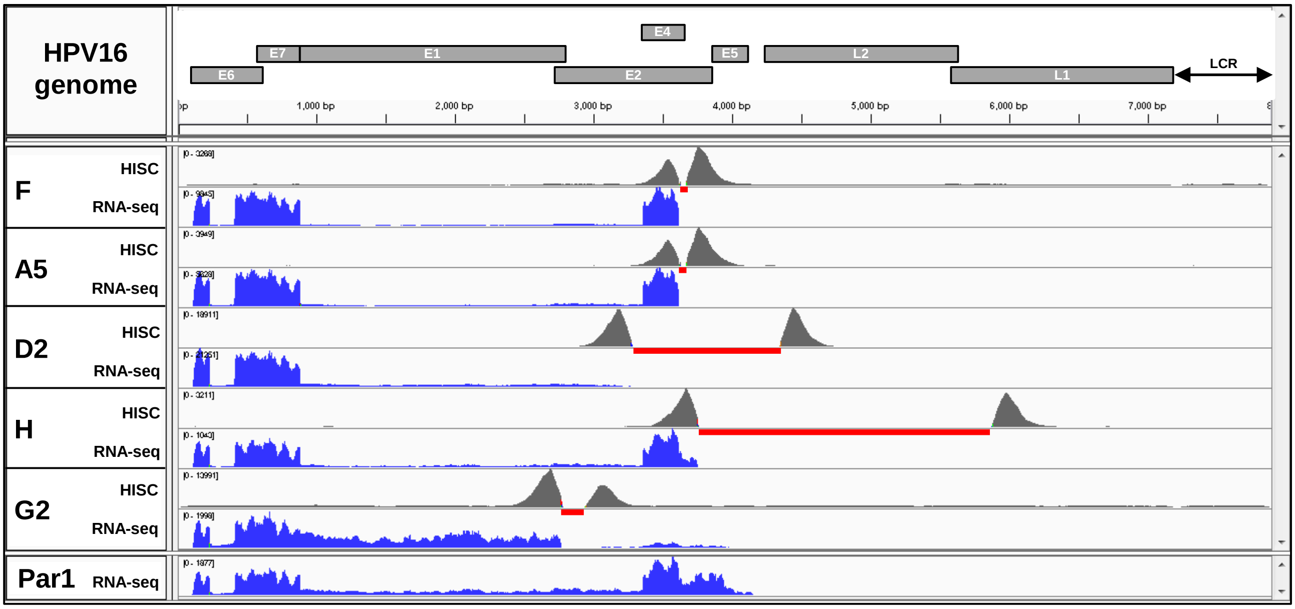


**Supplementary Figure 2. W12 integrant clone HPV16 genome length and breakpoint determination by Capture-ends and RNA-seq analysis.** Aligned to a cartoon of the HPV16 genome are DNA read peaks from Capture-ends analysis (grey peaks) determining breakpoints and deleted region of the HPV16 genome (red underline) as well as RNA read peaks (blue) denoting transcription across the virus genome for each of the W12 integrant clones used in the study along with RNA reads for the episomal W12 parental (Par1) polyclonal cell line for comparison.

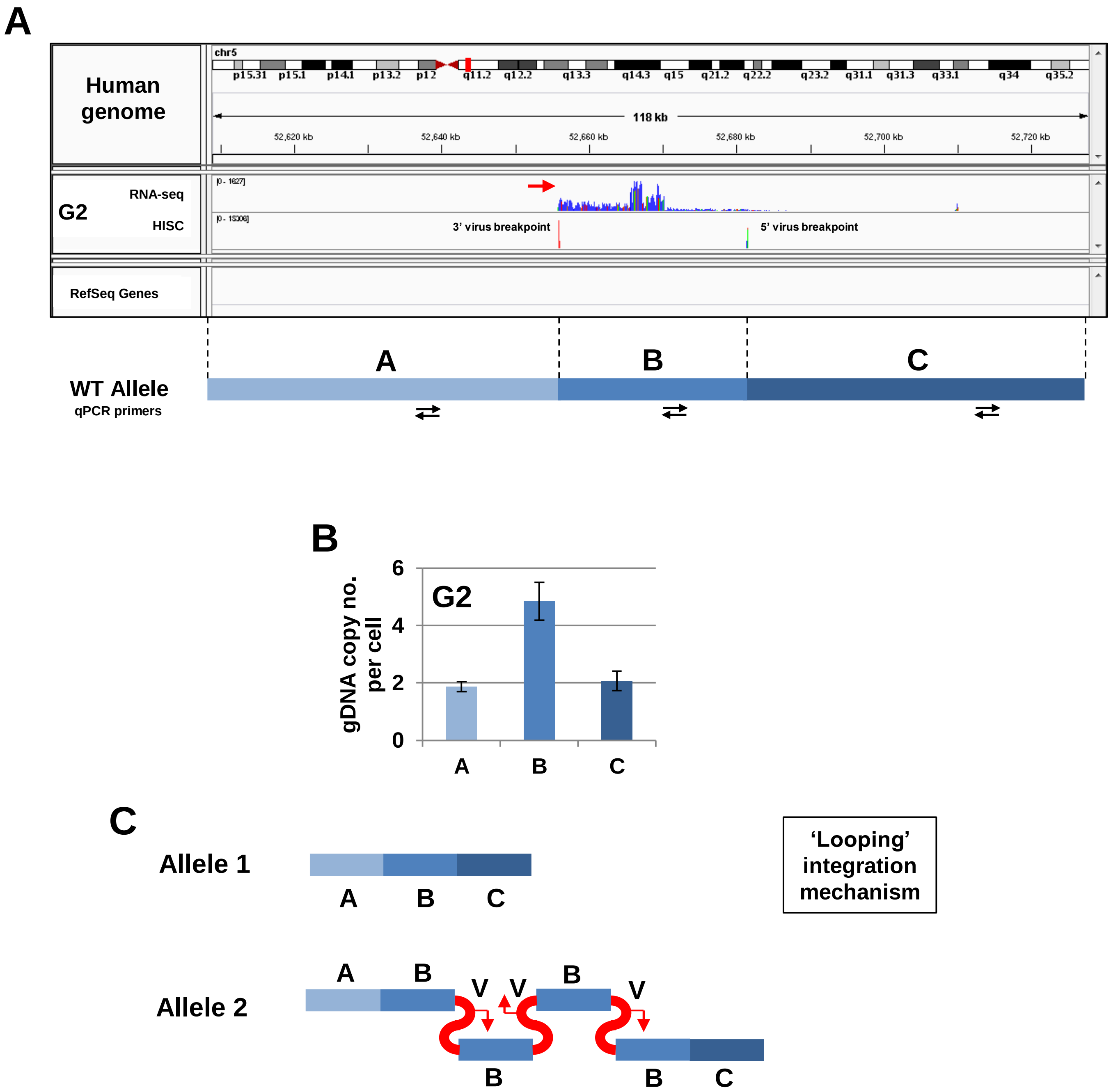


**Supplementary Figure 3.** Arrangement of gDNA at clone G2 integration site. (A) RNA-seq data (blue peaks) showing transcription from host sequences driven by the integrant HPV16 genome and HPV Integration Site Capture (HISC) data (red peaks) verifying virus-host breakpoints on the host genome (red arrow – direction of HPV16-host read-through transcription). Wild-type allele regions indicated below with approximate location of qPCR primer sites. (B) qPCR of genomic DNA regions to determine copy number after HPV16 genome integration in clone G2. (C) Determination of arrangement of gDNA sections after HPV16 genome integration through 'looping' mechanism, amplifying region B. Virus copy number (V, 3) based on Scarpini et al., 2014. Not to scale.

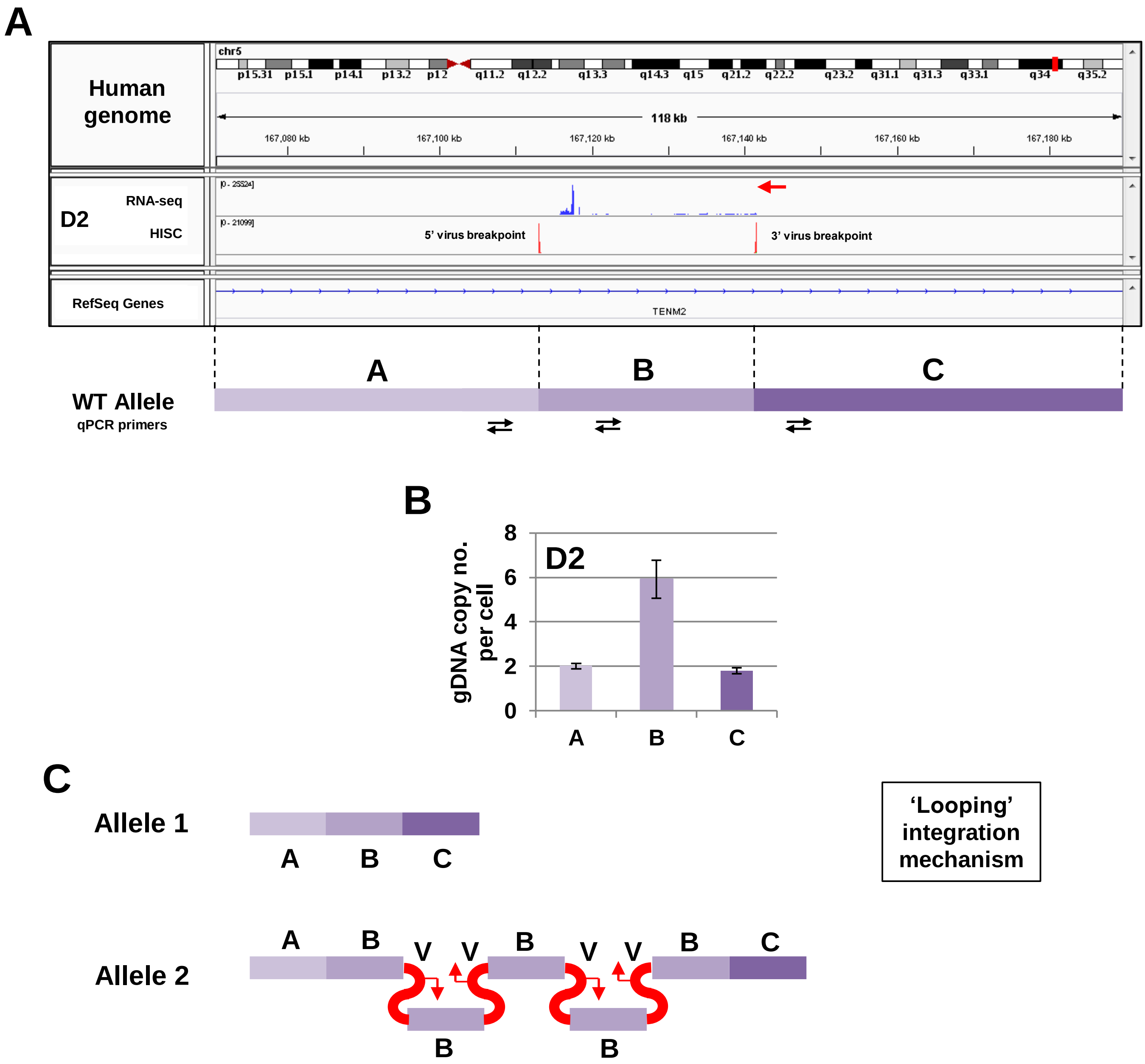


**Supplementary Figure 4. Arrangement of gDNA at clone D2 integration site.** (A) RNA-seq data (blue peaks) showing transcription from host sequences driven by the integrant HPV16 genome and HPV Integration Site Capture (HISC) data (red peaks) verifying virus-host breakpoints on the host genome (red arrow – direction of HPV16-host read-through transcription). Wild-type allele regions indicated below with approximate location of qPCR primer sites. (B) qPCR of genomic DNA regions to determine copy number after HPV16 genome integration in clone D2. (C) Determination of arrangement of gDNA sections after HPV16 genome integration through 'looping' mechanism, amplifying region B. Virus copy number (V, 4) from Scarpini et al., 2014. Not to scale.

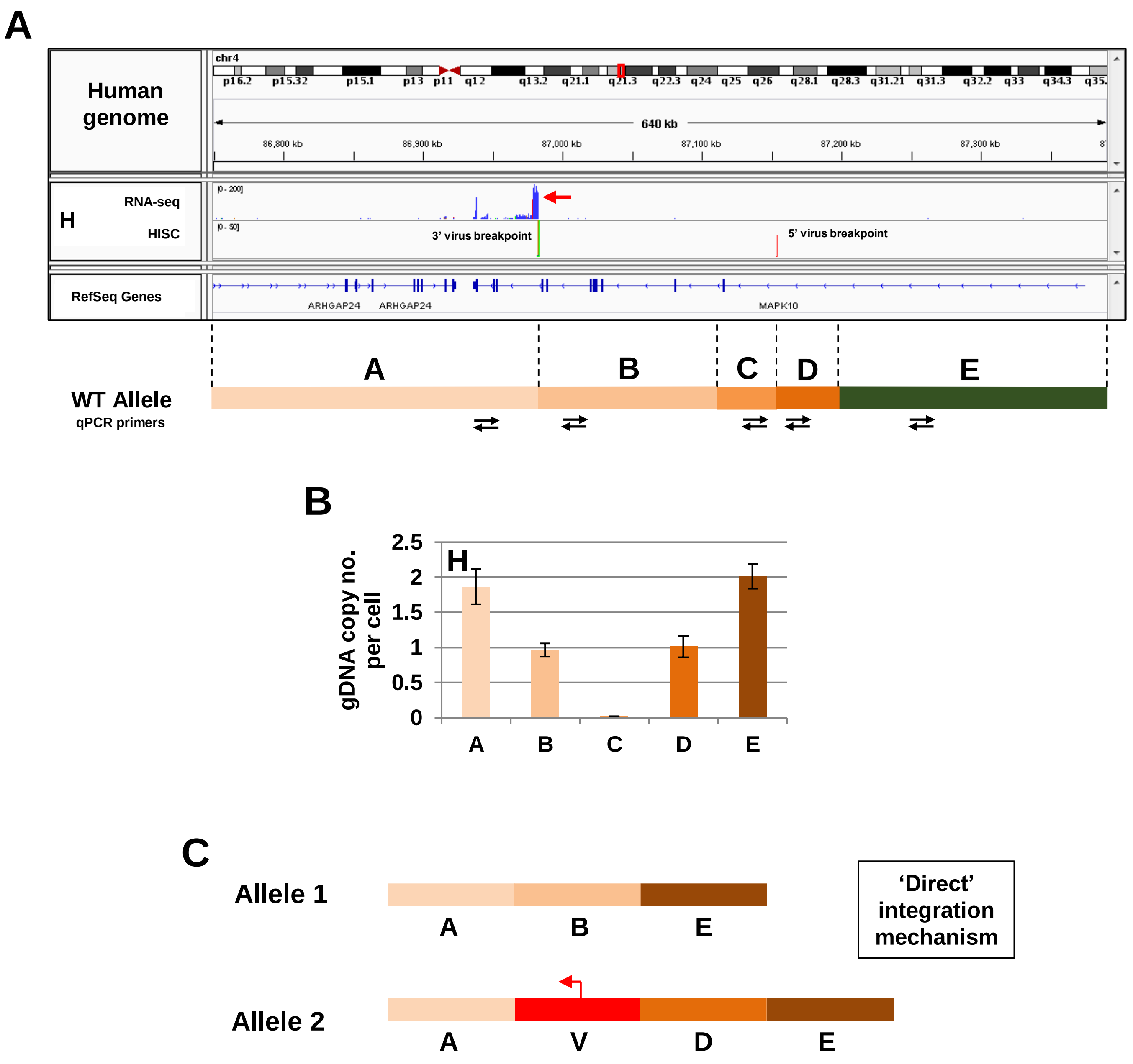


**Supplementary Figure 5. Arrangement of gDNA at clone H integration site.** (A) RNA-seq data (blue peaks) showing transcription from host sequences driven by the integrant HPV16 genome and HPV Integration Site Capture (HISC)data (red peaks) verifying virus-host breakpoints on the host genome (red arrow – direction of HPV16-host read-through transcription). Wild-type allele regions indicated below with approximate location of qPCR primer sites. (B) qPCR of genomic DNA regions to determine copy number after HPV16 genome integration in clone H. (C) Determination of arrangement of gDNA sections after HPV16 genome integration through 'direct' mechanism, causing deletion of reions including homozygous deletion of region C. Virus copy number (V, 1) from Scarpini et al., 2014. Not to scale. Target regions for qPCR based upon preliminary DNA-seq data not used in this study.

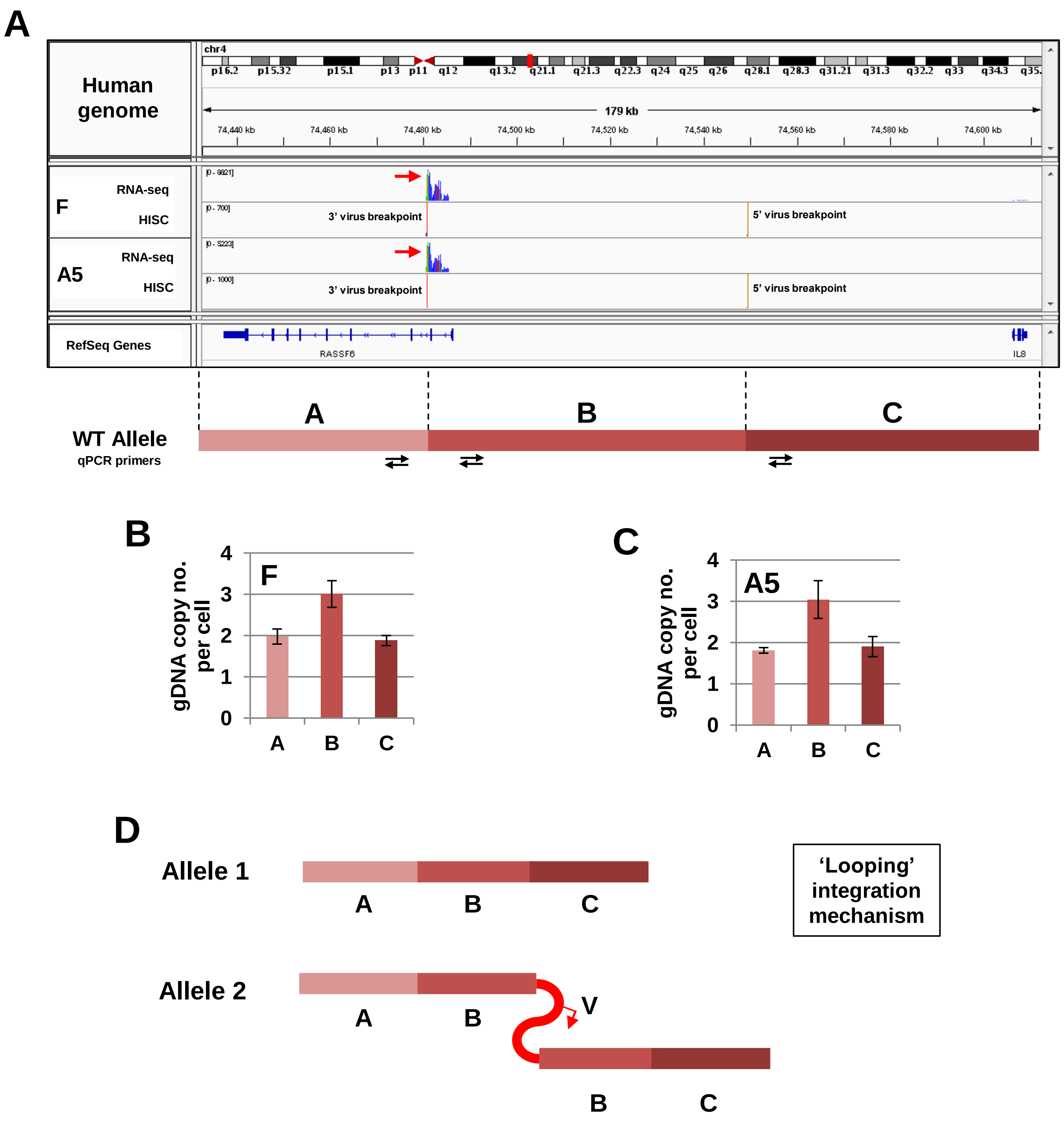


**Supplementary Figure 6. Arrangement of gDNA at clone F/A5 integration site.** (A) RNA-seq data (blue peaks) showing transcription from host sequences driven by the integrant HPV16 genome and HPV Integration Site Capture (HISC) data (red peaks) verifying virus-host breakpoints on the host genome (red arrow – direction of HPV16-host read-through transcription). Wild-type allele regions indicated below with approximate location of qPCR primer sites. (B) qPCR of genomic DNA regions to determine copy number after HPV16 genome integration in clone F and (C) A5. (D) Determination of arrangement of gDNA sections after HPV16 genome integration through 'looping' mechanism, amplifying region B. Virus copy number (V, 1) from Scarpini et al., 2014. Not to scale.

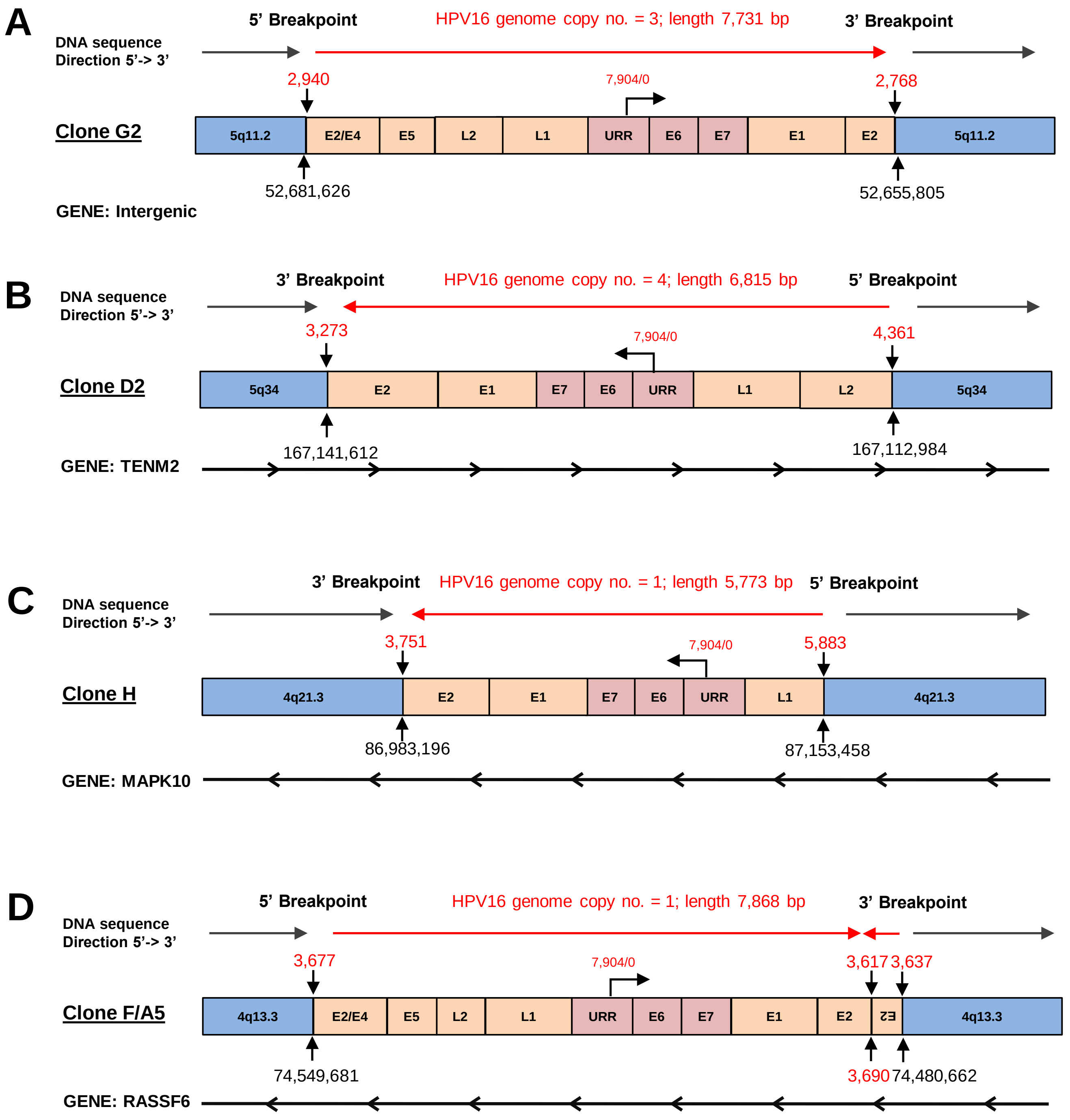


**Supplementary Figure 7. Summary of virus-host junction genomes at HPV16 integration sites.** In all schematics, host chromosomal DNA is shown in orange and the orientation indicated by the grey arrow above (5' to 3'). Integrated HPV16 DNA is shown in blue, with the viral oncogenes and LCR highlighted in red, and the direction of transcription from the viral early promoter shown by an arrow from the URR. The location of the viral breakpoint in base pairs is given above the junction, whereas the cellular DNA breakpoint in base pairs is given below the junction. The genome copy number and length of the integrated HPV16 genome is indicated in red above the schematic. When HPV16 has integrated into a host gene, the orientation is shown beneath the schematic in black. A) W12 clone G2, B) W12 clone D2, C) W12 clone H and D) W12 clones F/A5. (Virus genome copy number taken from Scarpini et al., 2014.)

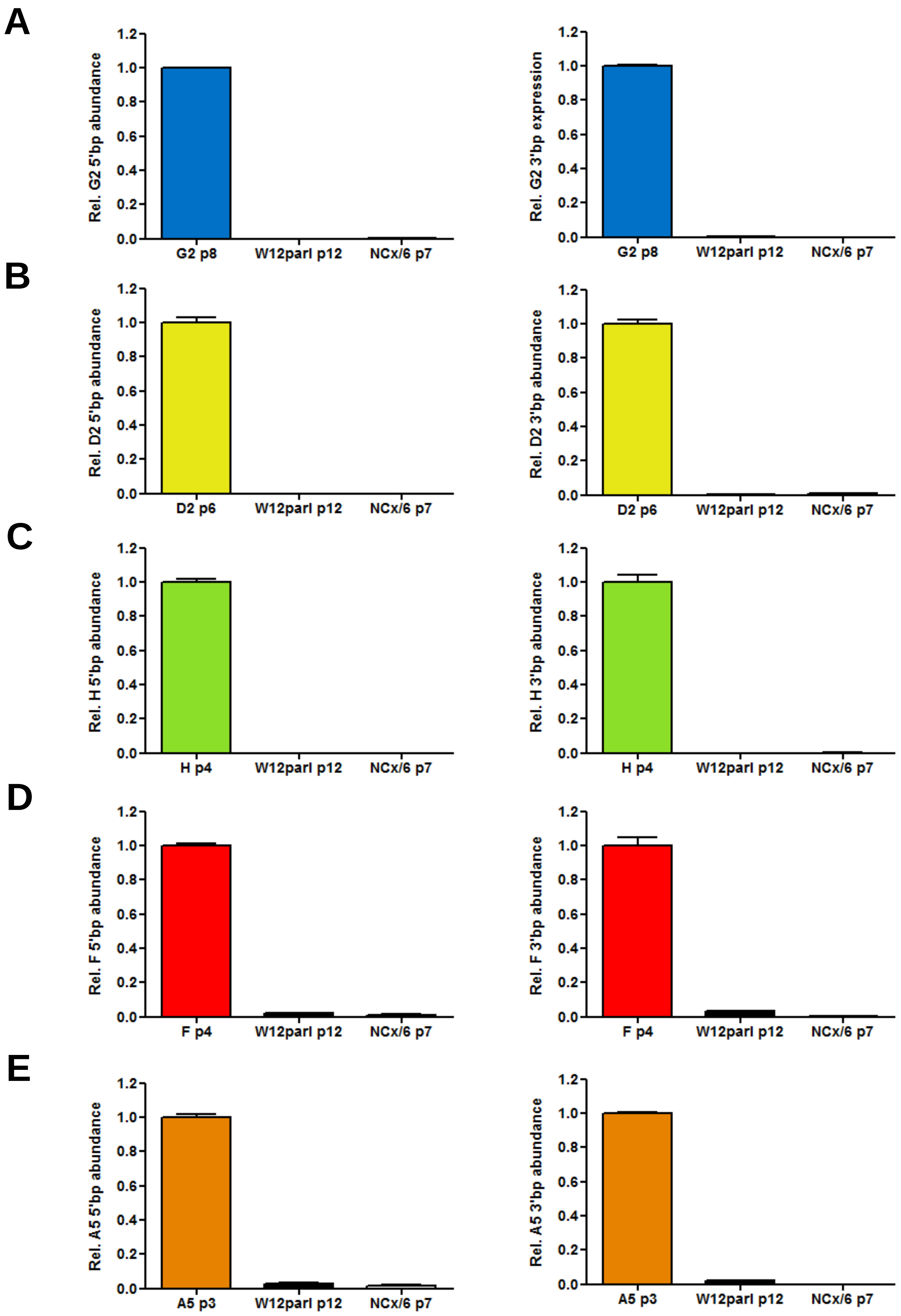


**Supplementary Figure 8. Confirmation of HPV16-host breakpoints by quantitative PCR.** To confirm the virus-host breakpoints found in each W12 integrant cloned line, pairs of qPCR primers were designed to amplify the 5' (left column) and 3' (right column) breakpoints from genomic DNA samples for clones (A) G2, (B) D2, (C) H, (D) F and (E) A5 in comparison to the episomal (W12par1 p12) cell line and HPV-negative cell line (NCx/6).

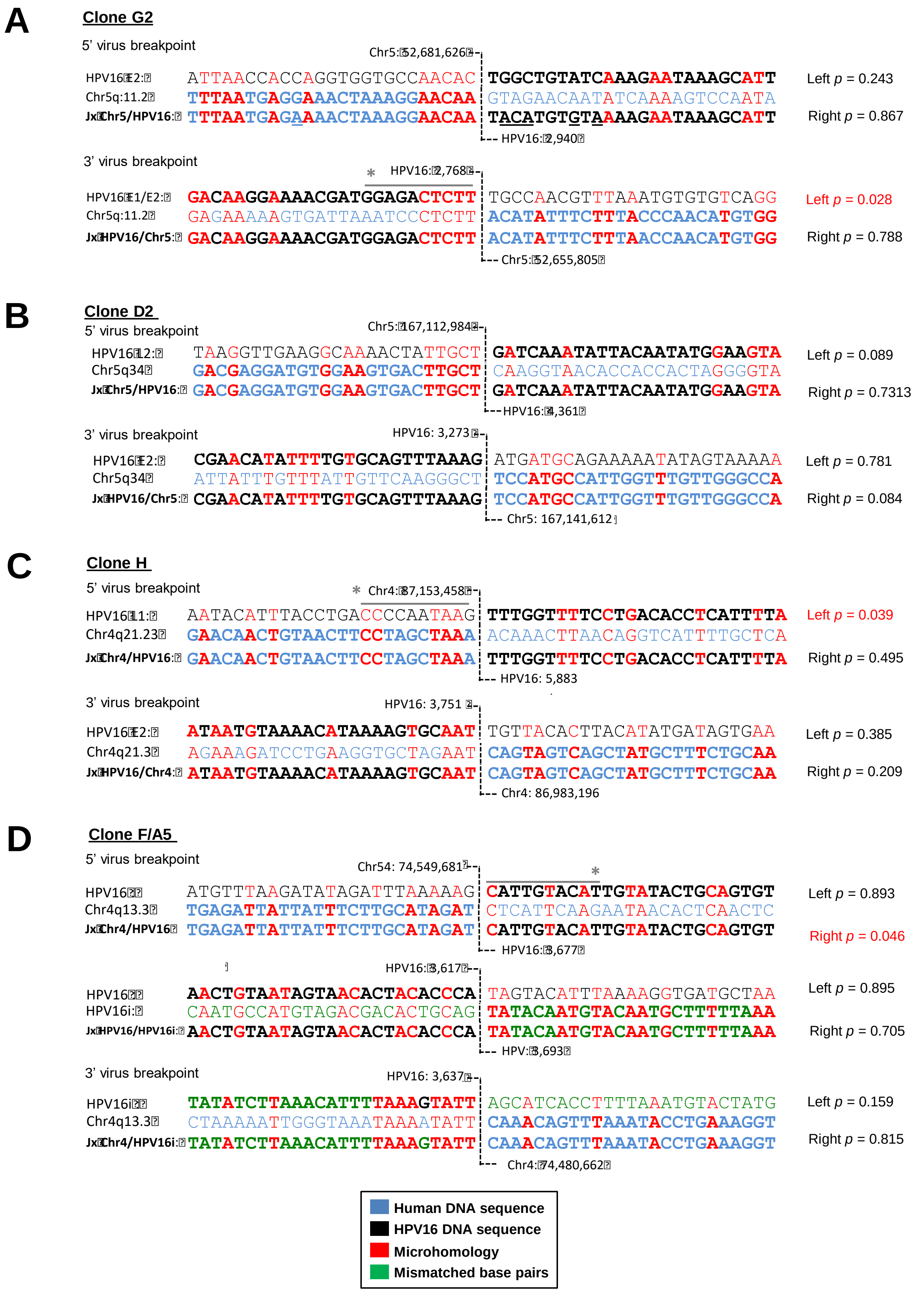


**Supplementary Figure 9. Regions of HPV16 and host sequence homology at the integration site.** Figures show comparisons between the virus-host sequences obtained by Sanger sequencing and the normal host and HPV16 genomic sequences, 25 nucleotides either side of the breakpoint (indicated by a central dotted line). HPV16 DNA sequence = black, inverted HPV16 DNA sequence = green, human DNA sequence = blue, homologous nucleotides = red. Significant levels of microhomology between host and HPV16 sequences were calculated by comparing the homology seen at the 10 nt directly either side of the breakpoint compared to 1000 nt of extended sequence which was shuffled 10,000 times and are indicated by a line above appropriate sequences. *$p < 0.05$ (highlighted in red).

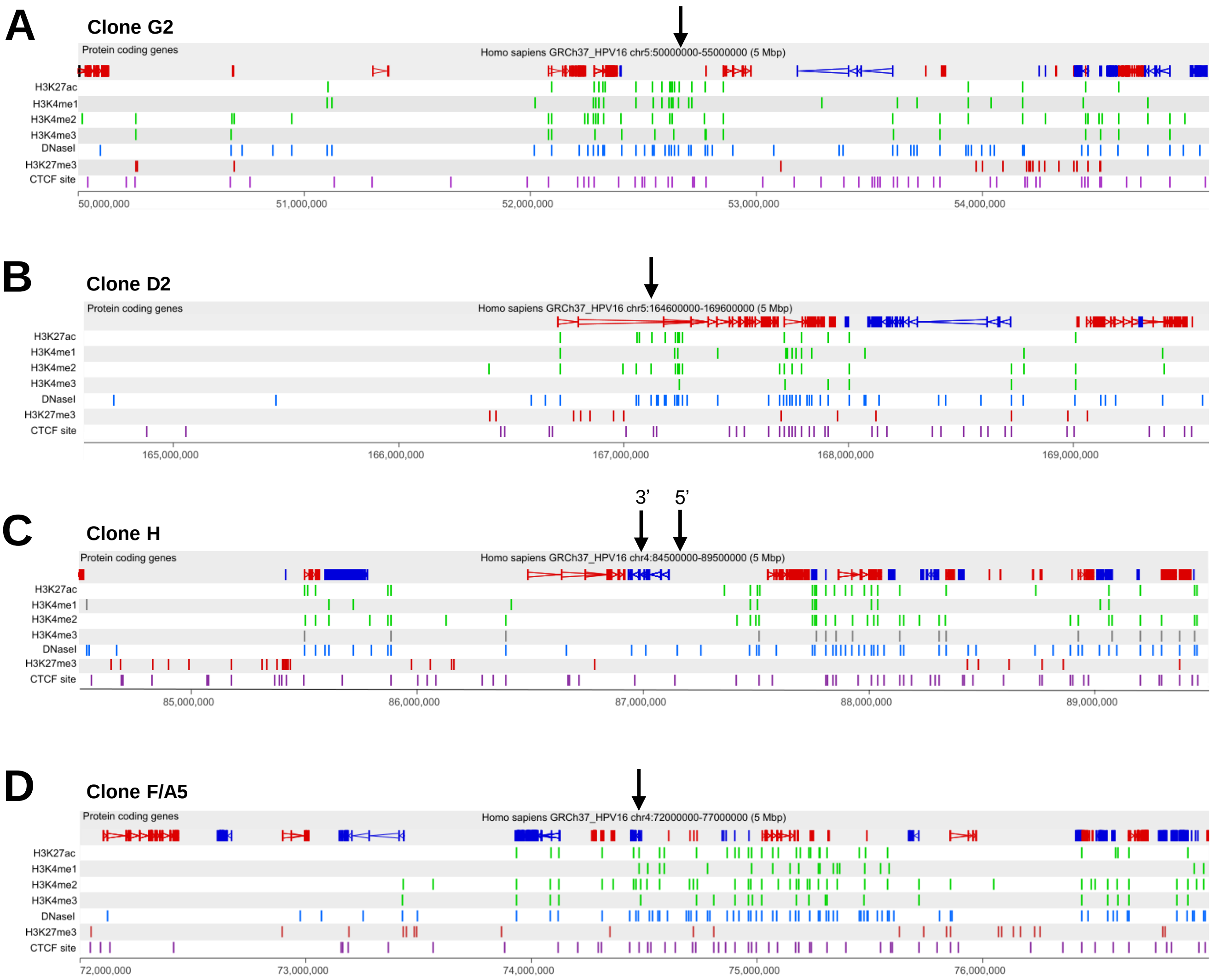


**Supplementary Figure 10. Analysis of host chromatin structure at the sites of HPV16 integration.** Each panel shows 5 Mb of the host genome across the integration loci for W12 clones (A) G2, (B) D2, (C) H, and (D) F/A5 with the virus integration site indicated by a black arrow. Protein coding genes are shown in the first track and the direction of each gene indicated by colour (red, forward; blue, reverse). ChIP-seq data from a normal human epidermal keratinocyte (NHEK) cell line is aligned with the host genome (taken from ENCODE). Post-translational histone modifications of active chromatin (H3K27ac, H3K4me1, H3K4me2, H3K4me3; green), repressive H3K27me3 (red), DNaseI hypersensitivity sites (blue) and CTCF sites (purple) are shown. Coordinates presented for each clone are indicated at the top of each figure. W12 clone H has both 5' and 3' ends of the HPV16 genome identified due to the length of the host genomic deletion.

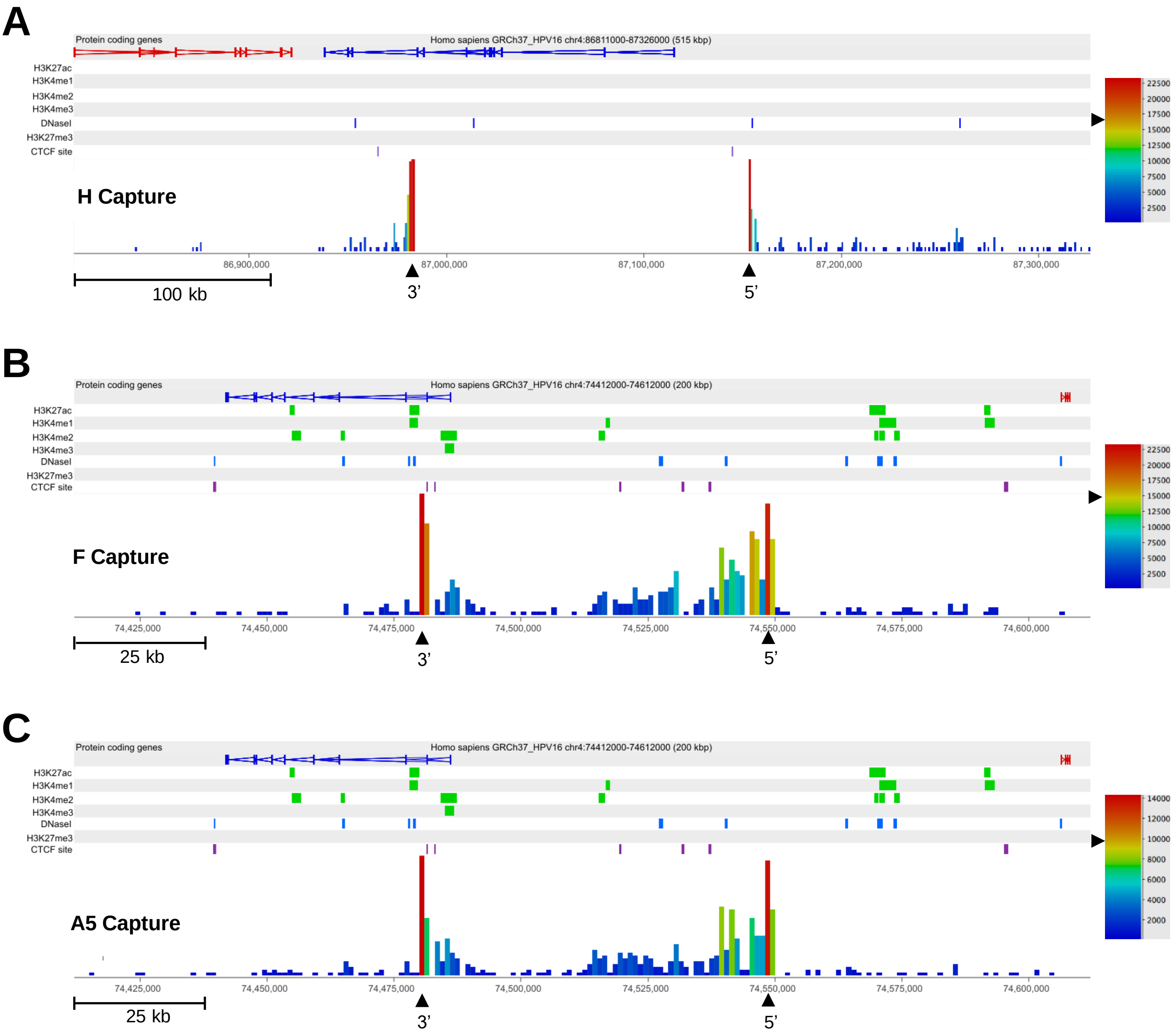


**Supplementary Figure 11. HPV16-host breakpoints identified in clones H, F and A5 by HPV16-specific Region Capture Hi-C.** (A) Capture Hi-C data is presented 515 kbp across the HPV16 integration locus for W12 clone H. The 5' and 3' breakpoints of the virus are indicated by the tallest red bars and are labelled with black arrowheads, running leftward due to the direction of virus sequence and without intermediate reads due to deletion of host sequence during 'direct' integration mechanism. (B) Capture Hi-C data is presented 200 kbp across the HPV16 integration locus for W12 clone F and (C) for W12 clone A5. The 5' and 3' breakpoints of the virus are indicated by the tallest red bars and are labelled with black arrowheads, being inverted in comparison to the direction of host sequence due to the 'looping' integration mechanism. In each panel, the scale bar represents the normalised read count. Additionally, protein-coding genes are shown in the first track, followed by the alignment of ChIP-seq data from the NHEK cell line (ENCODE). Post-translational histone modifications of active chromatin (H3K27ac, H3K4me1, H3K4me2, H3K4me3; green), repressive H3K27me3 (red), DNaseI hypersensitivity sites (blue) and CTCF sites (purple) are shown. Coordinates presented for each window are indicated at the top of each figure.

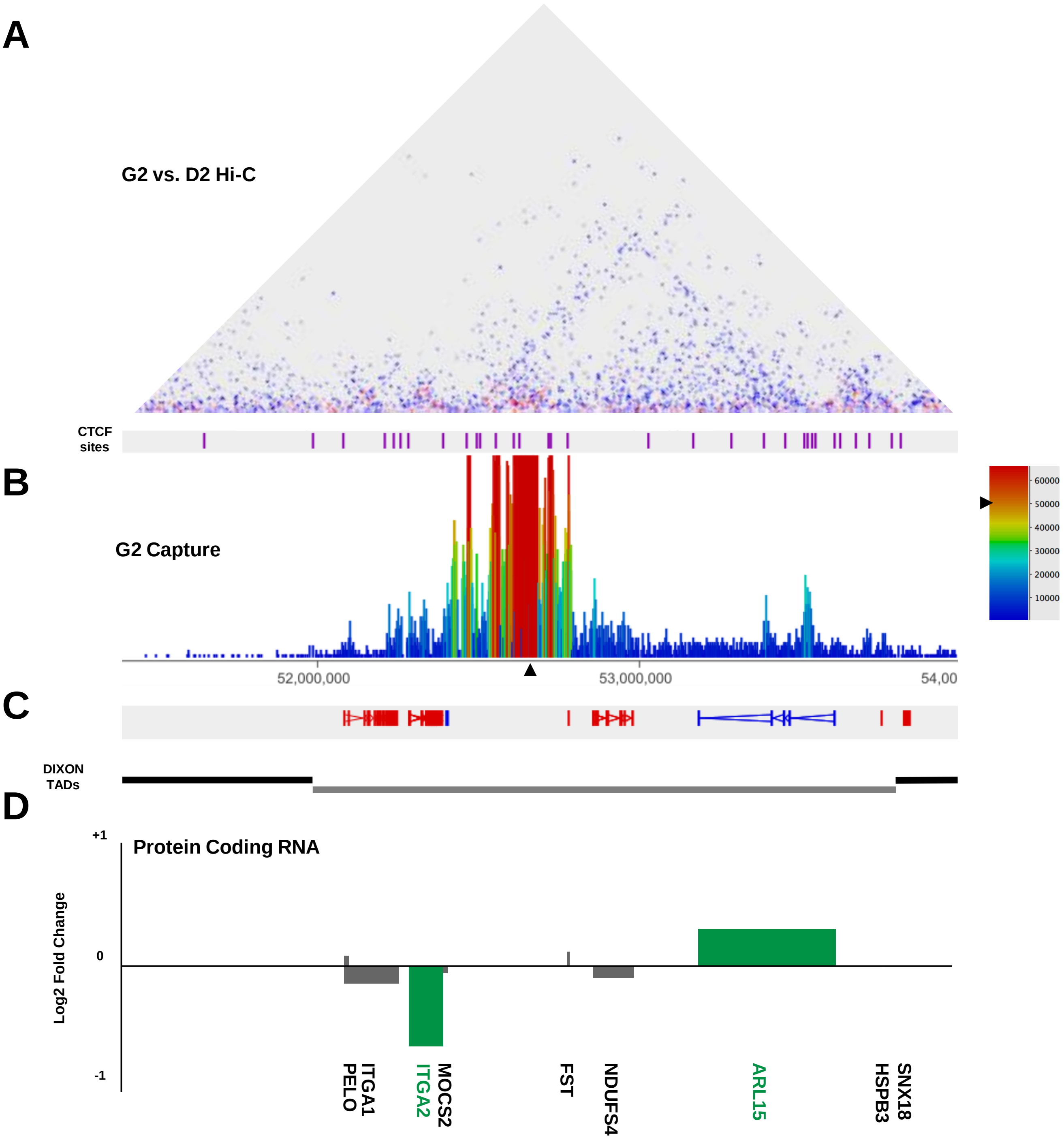


**Supplementary Figure 12. Chromosomal integration of HPV16 genomes causes host:host interaction changes within TADs.** (A) Comparative analysis of Hi-C libraries between clones G2 and D2 shows a decrease in a host:host interaction within the TAD of integration (blue triangle); aligned to sites of host CTCF interaction (purple lines). (B) Associated Capture Hi-C data is presented across Chr5: 51.5-54 Mbp. HPV16 integration site is indicated with a black arrow (scale bar represents the normalised read count). (C) Aligned protein coding genes (rightward, red; leftward, blue) and the extent of topologically associating domains (TADs) determined by Dixon et al. are shown below. (D) Chart indicating the transcript level of host protein coding genes within the 2.5 Mb region of W12 clone G2 relative to a 6-clone integrant average control level. All data is shown as a Log2 fold change with significant changes indicated by green bars. Gene length is indicated by width of the corresponding bar.

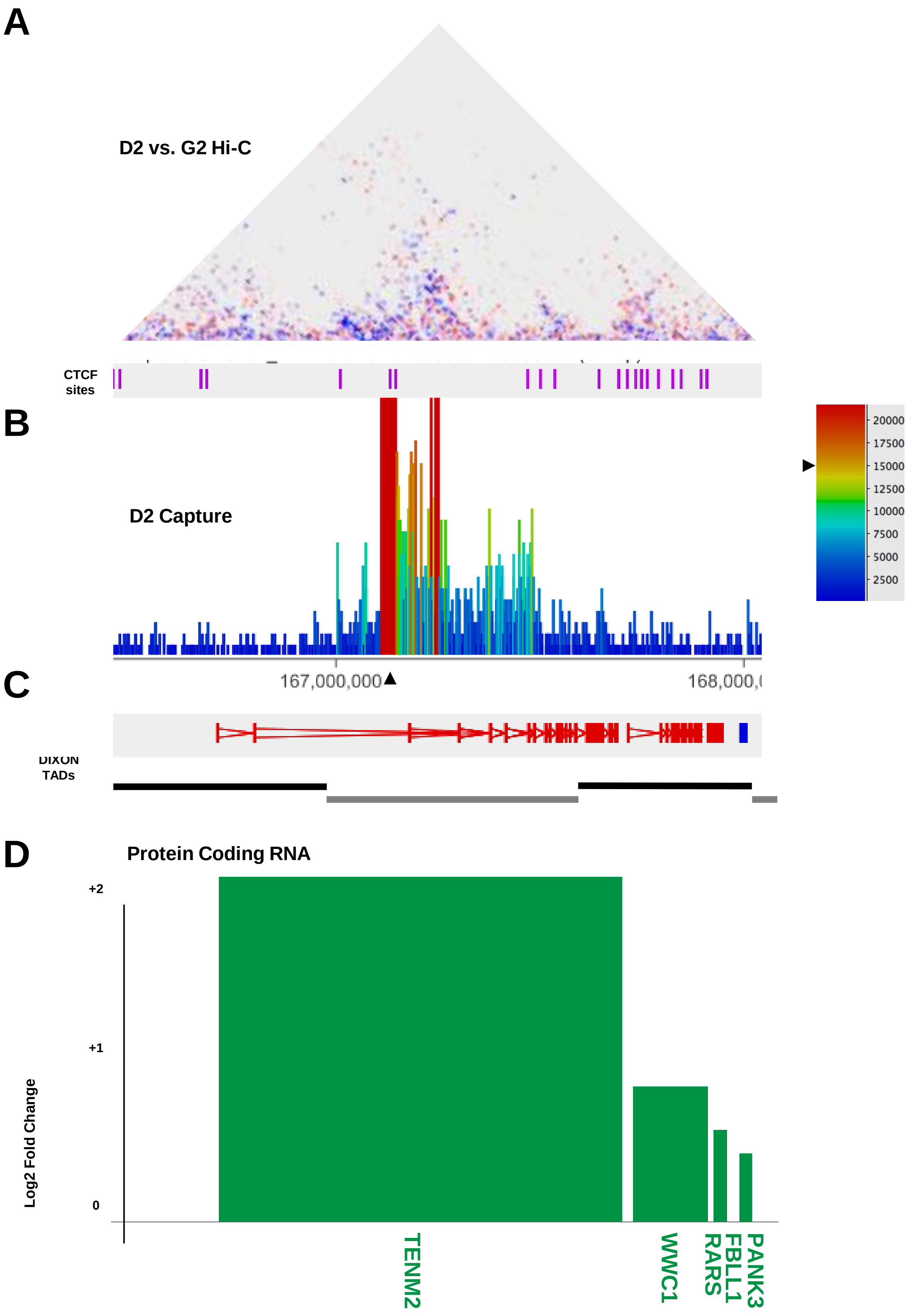


**Supplementary Figure 13. Chromosomal integration of HPV16 genomes causes transcript modulation without host:host interaction changes within TADs.** (A) Comparative analysis of Hi-C libraries between clones D2 and G2 shows no significant host:host interaction changes within the TAD of integration; aligned to sites of host CTCF interaction (purple lines). (B) Associated Capture Hi-C data is presented across Chr5: 166.5-168 Mbp. HPV16 integration site is indicated with a black arrow (scale bar represents the normalised read count). (C) Aligned protein coding genes (rightward, red; leftward, blue) and the extent of topologically associating domains (TADs) determined by Dixon et al. are shown below. (D) Chart indicating the transcript level of host protein coding genes within the 1.5 Mb region of W12 clone D2 relative to a 6-clone integrant average control level. All data is shown as a Log2 fold change with significant changes indicated by green bars. Gene length is indicated by width of the corresponding bar.

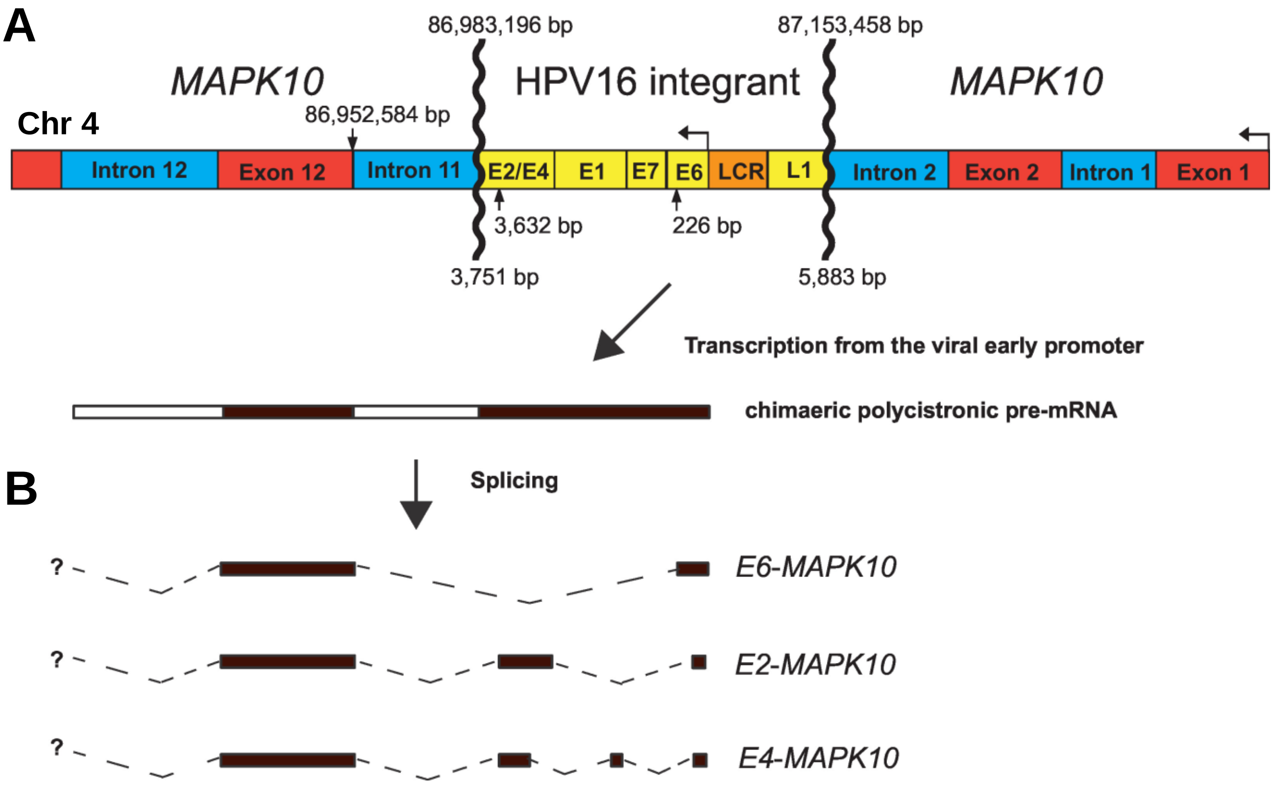


**Supplementary Figure 14. Integration of HPV16 genomes into host chromosomes in W12 clone H causes virus:host spliced fusion transcripts.** (A) Diagram summarises HPV16:host breakpoints and deletion of introns/exons in *MAPK10* gene with (B) spliced fusion transcripts found by RNA-sequencing in W12 clone H.

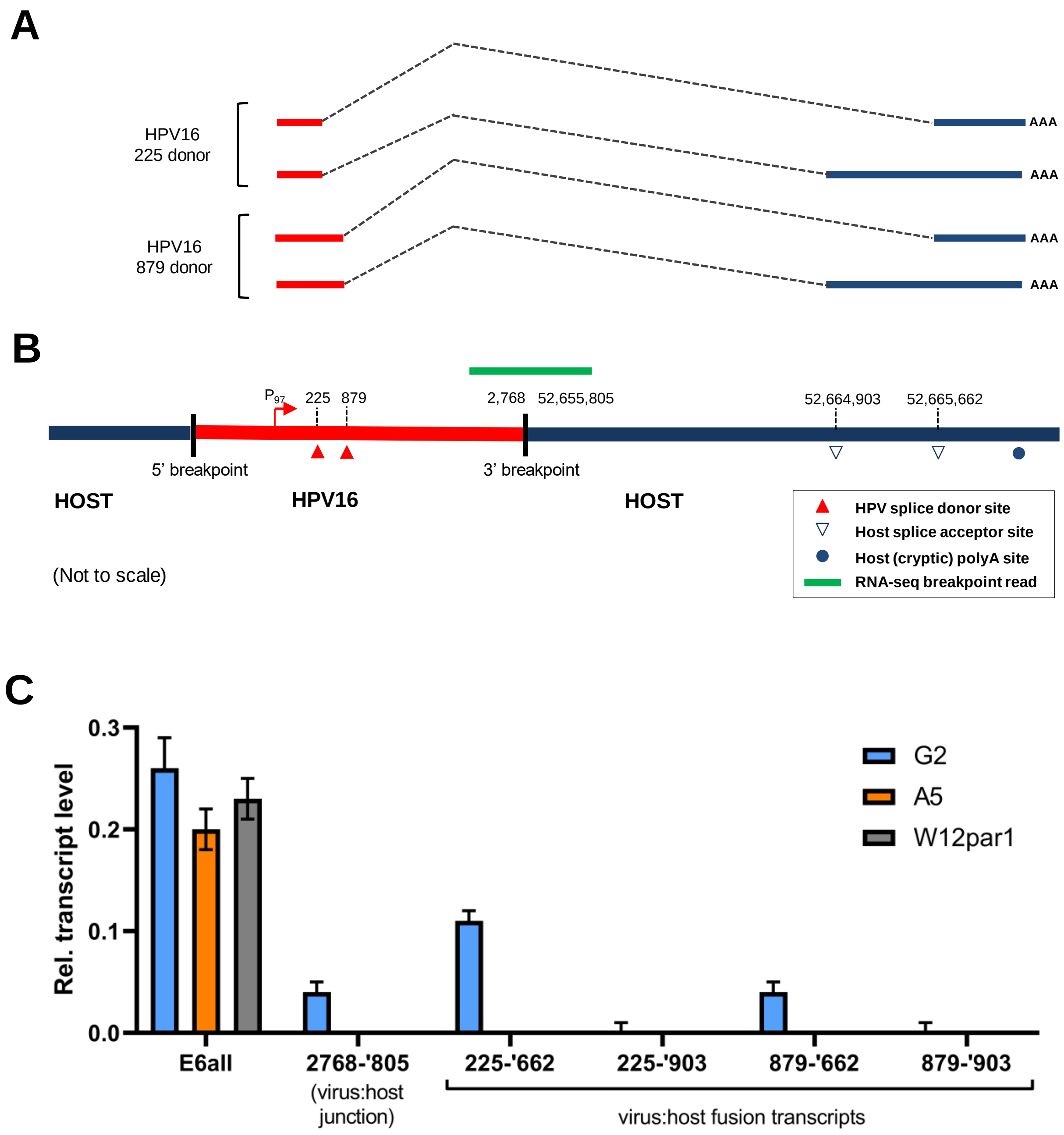


**Supplementary Figure 15. Integration of HPV16 genomes into host chromosomes in W12 clone G2 causes virus:host spliced fusion transcripts.** (A) Diagram summarises HPV16:host fusion transcripts found by RNA-sequencing in W12 clone G2. (B) Mapping of the fusion transcript HPV16 splice donor locations and host splice acceptor sites, (C) Determination of G2-specific fusion transcript expression levels via qPCR in comparison to total E6 coding transcripts (E6all) in clones G2 (blue), A5 (orange) and episomal W12par1 (grey).

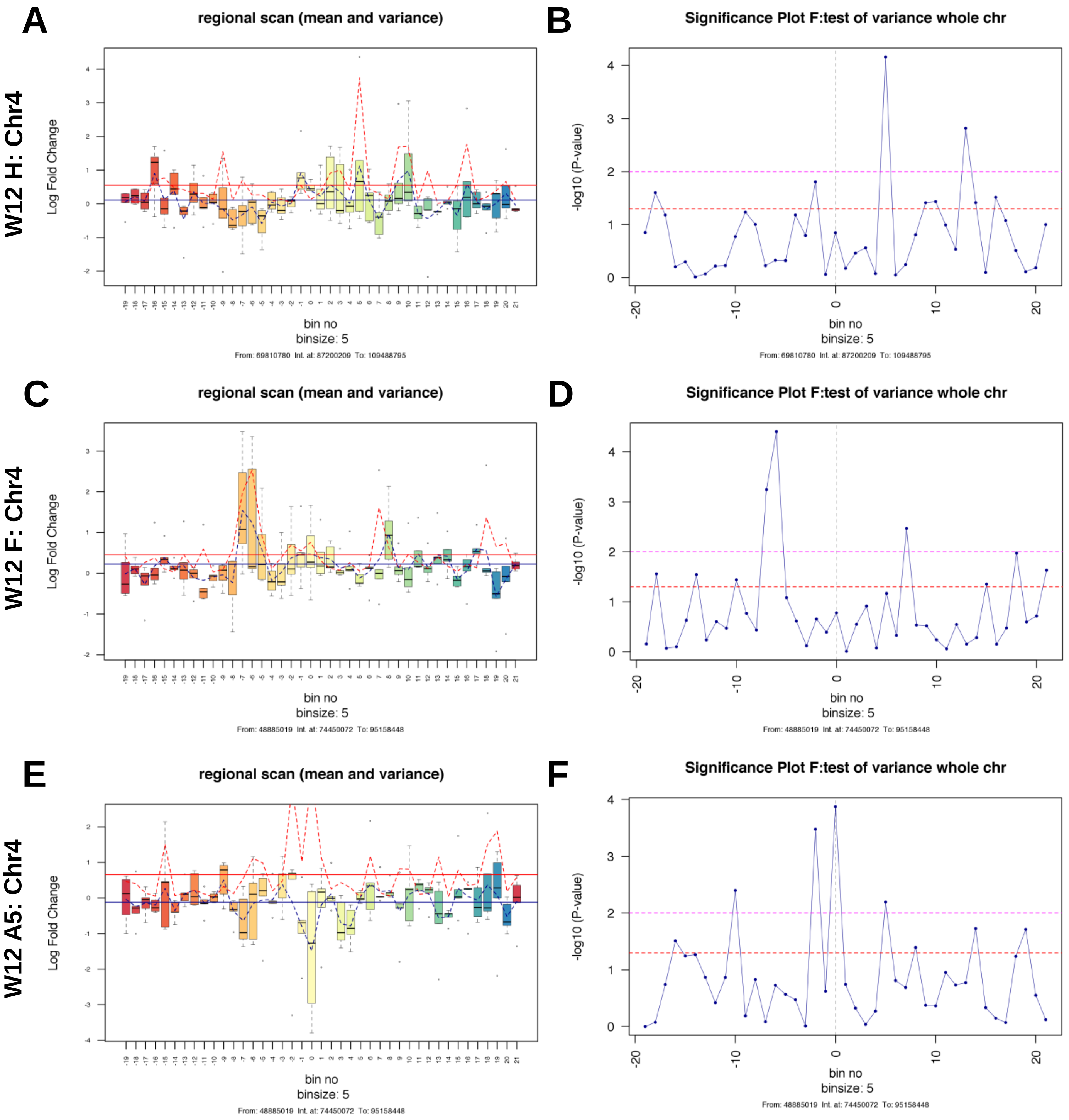


**Supplementary Figure 16. Variance in host gene expression across the host genomic region containing the HPV16 integration site in W12 clones H, F and A5.** Each left panel indicates the range and variance of host gene expression in W12 integrant clones [A) W12 H, C) W12 F and E) W12 A5], focussing on 100 genes either side of the HPV16 integration site. For each clone, gene expression levels were compared with a 6-clone integrant average control level. In each panel, the HPV16 integration site is centred on 'bin 0'. Each bin contains five genes, with no overlap between bins. The box and whisker plots illustrate the range of gene expression levels within each bin, with the bar indicating median values, the box the IQR and the whiskers the range. The mean gene expression across the whole chromosome is indicated by the solid blue line, while the mean level of gene expression across individual bins is shown by the dotted blue line. The mean variance of gene expression across the whole chromosome is indicated by the solid red line, while the mean level of gene expression across individual bins is shown by a dotted red line. Each right hand panel shows the significance of the variance in gene expression within each bin [B) W12 H, D) W12 F and F) W12 A5]. Each point represents a five-gene bin, corresponding to those in the left-hand panels. The horizontal lines indicate the significance of the variance in each bin, compared with the variance in gene expression across the whole chromosome (above the dashed red line, $p<0.05$; above the dashed pink line, $p<0.01$).

### Supplementary Table 1. RNA-seq fusion reads including HPV16 sequence from W12 cells.

| Target | Host chromosome breakpoint or HPV16 breakpoint (nt) | HPV16 breakpoint (nt) | No. of reads spanning fusion (n=2)* | Splice or HPV16:host junction (3' end) | Verified (V) or not verified (N) |
|---|---|---|---|---|---|
| **A5** | Chr4: 74,480,662 | 3,637 | 1402 + 1756 | Junction | V |
| | HPV16: 3,617 | 3,690 | 1619 + 1959 | Junction | V |
| **F** | Chr4: 74,480,662 | 3,637 | 2013 + 436 | Junction | V |
| | HPV16: 3,617 | 3,690 | 2342 + 2066 | Junction | V |
| **G2** | Chr5: 52,655,805 | 2768 | 375 + 414 | Junction | V |
| | Chr5: 52,664,903 | 225 | Not found† | Splice | V |
| | | 879 | 14* + 11* | Splice | V |
| | Chr5: 52,665,662 | 225 | 119 + 92* | Splice | V |
| | | 879 | 306 +265 | Splice | V |
| **H** | Chr4: 86,952,584 | 225 | 16* + 19* | Splice | N |
| | | 879 | 17* + 16* | Splice | N |
| | Chr4: 86,983,196 | 3,751 | 125 + 153 | Junction | V |
| **D2** | Chr5: 167,117,611 | 879 | 624 + 635 | Splice | N |
| | Chr5: 167,118,361 | 225 | 3598 + 3826 | Splice | N |
| | | 879 | 8245 + 8854 | Splice | N |
| | Chr5: 167,120,631 | 225 | 115 + 98* | Splice | N |
| | | 879 | 191 +184 | Splice | N |
| | Chr5: 167,122,165 | 879 | 349 + 365 | Splice | N |
| | Chr5: 167,127,882 | 879 | 106 + 123 | Splice | N |
| | Chr5: 167,135,217 | 225 | 104 + 134 | Splice | N |
| | | 879 | 606 + 737 | Splice | N |
| | Chr5: 167,141,282 | 879 | 137 + 155 | Splice | N |
| | Chr5: 167,141,612 | 3273 | 814 + 929 | Junction | V |
| **Par1** | Not found | Not found | N/A | N/A | N/A |
| **NCx/6** | Not found | Not found | N/A | N/A | N/A |

*cut off for fusion reads >100, unless otherwise indicated.
†splice was found by PCR and Sanger sequencing.

### Supplementary Table 2. Primers for PCR amplification and Sanger sequencing of HPV16-host breakpoints.

| Target | Forward primer (5' to 3') | | Reverse sequence (5' to 3') | |
|---|---|---|---|---|
| | Species | | Species | |
| **F/A5 3' breakpoint** | HPV16 | AAGGGCCCTAGCAGGTTTTA | Host | CACCGAAGAAACACAGACGA |
| **F/A5 5' breakpoint** | Host | TGGTCACGTTGCCATTGACT | HPV16 | GGGCAGTGTGGCAGTAGTTA |
| **H 3' breakpoint** | HPV | GCACCGAAGAAACACAGACG | Host | TCCTTCCCTCCCTAACAGCAT |
| **H 5' breakpoint** | Host | TGGGTCAGTGGTTTGATTGA | HPV | TGGGGATCCTTTGCCCCAGTGT |
| **D2 3' breakpoint** | HPV16 | ACTGTGGTAGAGGGTCAAGT | Host | GGGGAAGGTGGCATCTCTTA |
| **D2 5' breakpoint** | Host | CTTTGCCACGGGACAAGTAT | HPV16 | GGATCGGAAGGGCCCACAGGA |
| **G2 3' Breakpoint** | HPV16 | CAGTGCCCTGTTGGAACTACA | Host | GCTGTTGACCTCTTTGGGGT |
| **G2 5' breakpoint** | Host | TTACTCATGCCACCACACCT | HPV16 | CAACTTGACCCTCTACCACAGT |

### Supplementary Table 3. Primer pairs for PCR and qPCR amplification of clone G2 HPV16:host spliced RNAs.

| Target | Forward (HPV16) Primer (5' to 3') | Reverse (Host) Primer (5' to 3') |
|---|---|---|
| **G2 splice HPV16:225 to Chr5:52,664,903** | GCAACAGTTACTGCGACGTG | TGCTGTTGATTGGTCCTCCA |
| **G2 splice HPV16:879 to Chr5:52,664,903** | GAACCGGACAGAGCCCATTA | TGCTGTTGATTGGTCCTCCA |
| **G2 splice HPV16:225 to Chr5:52,665,662** | GCAACAGTTACTGCGACGTG | ATTCTCGTGGGAGGGAAAGC |
| **G2 splice HPV16:879 to Chr5:52,665,662** | GAACCGGACAGAGCCCATTA | ATTCTCGTGGGAGGGAAAGC |
| **E6all (Scarpini et al., 2014)** | TGTTTCAGGACCCACAGGAGC | CGCAGTAACTGTTGCTTGCAG |

### Supplementary Table 4. Primers for qPCR amplification of HPV16-host breakpoints.

| Target | Forward Primer (5' to 3') | | Reverse Primer (5' to 3') | |
|---|---|---|---|---|
| | Species | | Species | |
| **G2 5' breakpoint** | Human | ACCACACCTGGCTGAGAAAA | HPV16 | AGTTGCAGTTCAATTGCTTGT |
| **G2 3' breakpoint** | HPV16 | AAGTTTGCACGAGGACGAGG | Human | TCATCTGTGTTTTGAGCCACAT |
| **D2 5' breakpoint** | Human | TGACGAGGATGTGGAAGTGAC | HPV16 | ACCCGACCCTGTTCCAATTC |
| **D2 3' breakpoint** | HPV16 | TGTTCATGAAGGGATACGAACA | Human | CCCAAGTCCATTGAATCCTG |
| **H 5' breakpoint** | HPV16 | TGCAAAGATGTTTTAATGTCCCA | Human | ACCAGCCGCTGTGTATCTG |
| **H 3' breakpoint** | Human | TGCAGAAAGCATAGCTGACTACT | HPV16 | ACTGCAGTGTCGTCTACATGG |
| **F/A5 5' breakpoint** | Human | CTCTGCCTGCACATGACTTG | HPV16 | TGTCCAATGCCATGTAGACG |
| **F/A5 3' breakpoint** | HPV16 | CAGCTCACACAAAGGACGGA | Human | TGGGACTTTTACCAAAGCATGT |
| **GAPDH** | Human | CGGCTACTAGCGGTTTTACG | Human | AAGAAGATGCGGCTGACTGT |

### Supplementary Table 5. Primers for qPCR amplification of host gDNA.

| Target | Forward Primer (5' to 3') | Reverse Primer (5' to 3') |
|---|---|---|
| **G2: A** | TCGCGTACTTGGCTTTCAGT | AACCACACAACCTGCACTCA |
| **G2: B** | TGTGTAGCTCAGGCACATGG | AGCTGCGAGATCTAATGGGC |
| **G2: C** | AGAGGATTGGCTGGGTGTTG | TCCCCATTGTGCTGCTTGAT |
| **D2: A** | CTCCATGTGGATGTGGCAGT | GAAGAGAAGCTGGCACTGGT |
| **D2: B** | CACACTTGCCTGGTTCTTGC | GTTCCAGATGGAGCAGCCTT |
| **D2: C** | ACTCCTGTCACGAAGGCTTG | AGCAGATTGTGGGAGAAGGC |
| **H: A** | ACCGCCCCTTCTACCTACAT | GTTCCCCATCTCCCTCTCCT |
| **H: B** | TGGGCAGAAGCAAGAGTCAG | TCTGCCACGAGGAATCACAC |
| **H: C** | CCAGCATGCCATTGGCTTTT | CACTGCCTTGTTTGCTGCAT |
| **H: D** | GGAATGGAGGCCCCAAAAGA | GCATGTGCAGACAAAGCACA |
| **H: E** | TGCAATGGCTTCCCTGAGAG | TGGCAAGCCATCTCACCATT |
| **F/A5: A** | TGGGCACACAGACACAACAT | ACCAAGTTGCCCAGACCAAA |
| **F/A5: B** | ACTGCCCTCAGTTCAAGTGG | GAGGGCACCTTGAGCAGAAT |
| **F/A5: C** | TCTGCCATGTACCCCTGAGA | GTGCCTCATTTCAGTCCCCA |
| **TLR2** | GGCCAGCAAATTACCTGTGTG | AGGCGGACATCCTGAACCT |
| **IFNβ** | TTGAATGGGAGGCTTGAATACTG | AATGCGGCGTCCTCCTTCT |

### Supplementary Table 6. MboI restriction sites and RNA baits for HPV16-specific genome cleavage and capture.

| | MboI restriction fragments in HPV16 genome | | | RNA baits | | | | |
|---|---|---|---|---|---|---|---|---|
| **#** | **Ends** | **Coordinates** | **Length (bp)** | **HPV16 gene** | **Coordinates** | **Direction** | **Length (bp)** | **gBlock** |
| **1** | MboI-MboI | 525-621 | 97 | E6/E7 | 529-622 | F | 93 | 1 |
| **2** | MboI-MboI | 622-870 | 249 | E7 | 624-744 | F | 120 | 1 |
| | | | | E7/E1 | 872-752 | R | 120 | 1 |
| **3** | MboI-MboI | 871-3479 | 2609 | E1 | 874-933 | F | 59 | 1 |
| | | | | E2 | 3480-3361 | R | 119 | 1 |
| **4** | MboI-MboI | 3480-4360 | 881 | E2 | 3482-3602 | F | 120 | 1 |
| | | | | L2 | 4363-4242 | R | 121 | 1 |
| **5** | MboI-MboI | 4361-4519 | 159 | L2 | 4364-4483 | F | 119 | 1 |
| **6** | MboI-MboI | 4520-4537 | 18 | | | | | |
| **7** | MboI-MboI | 4538-5071 | 534 | L2 | 4541-4660 | F | 119 | 1 |
| | | | | L2 | 5073-4953 | R | 120 | 2 |
| **8** | MboI-MboI | 5072-6150 | 1079 | L2 | 5075-5194 | F | 119 | 2 |
| | | | | L1 | 6150-6030 | R | 120 | 2 |
| **9** | MboI-MboI | 6151-6950 | 800 | L1 | 6152-6271 | F | 119 | 2 |
| | | | | L1 | 6950-6830 | R | 120 | 2 |
| **10** | MboI-MboI | 6951-7013 | 63 | | | | | 2 |
| **11** | MboI-MboI | 7014-524 | 1415 | L1 | 7014-7134 | F | 120 | 2 |
| | | | | E6 | 525-406 | R | 119 | 2 |
